# FOXP3^+^ regulatory T cells use heparanase to access IL-2 bound to ECM in inflamed tissues

**DOI:** 10.1101/2023.02.26.529772

**Authors:** Hunter A. Martinez, Ievgen Koliesnik, Gernot Kaber, Jacqueline K. Reid, Nadine Nagy, Graham Barlow, Ben A. Falk, Carlos O. Medina, Aviv Hargil, Israel Vlodavsky, Jin-Ping Li, Magdiel Pérez-Cruz, Sai-Wen Tang, Everett H. Meyer, Lucile E. Wrenshall, James D. Lord, K. Christopher Garcia, Theo D. Palmer, Lawrence Steinman, Gerald T. Nepom, Thomas N. Wight, Paul L. Bollyky, Hedwich F. Kuipers

## Abstract

FOXP3^+^ regulatory T cells (Treg) depend on exogenous IL-2 for their survival and function, but circulating levels of IL-2 are low, making it unclear how Treg access this critical resource *in vivo*. Here, we show that Treg use heparanase (HPSE) to access IL-2 sequestered by heparan sulfate (HS) within the extracellular matrix (ECM) of inflamed central nervous system tissue. HPSE expression distinguishes human and murine Treg from conventional T cells and is regulated by the availability of IL-2. HPSE^-/-^ Treg have impaired stability and function *in vivo*, including the experimental autoimmune encephalomyelitis (EAE) mouse model of multiple sclerosis. Conversely, endowing Treg with HPSE enhances their ability to access HS-sequestered IL-2 and their tolerogenic function *in vivo*. Together, these data identify novel roles for HPSE and the ECM in immune tolerance, providing new avenues for improving Treg-based therapy of autoimmunity.

**One-Sentence Summary:** Regulatory T cells use heparanase to strip IL-2 bound to extracellular matrix within inflamed tissues, thereby supporting their homeostasis and function.

## Main

In healthy individuals, immune tolerance is maintained by populations of regulatory T cells, including FOXP3^+^ regulatory T cells (Treg)^1^. However, in patients with autoimmune diseases such as multiple sclerosis (MS), Treg fail to control the autoreactive immune response central to their pathology. Although the frequency of circulating Treg is not different in MS patients compared to healthy controls, patient-derived Treg exhibit loss of *in vitro* function, as well as impaired FOXP3 expression levels^2–5^. Moreover, numbers of Treg within MS lesions have been reported to be low or non-existing, despite the presence of CD4^+^ T cell infiltrates^6^. This suggests that factors that regulate local Treg homeostasis and function may contribute to the loss of tolerance central to autoimmune conditions.

One factor that governs Treg homeostasis is the cytokine interleukin 2 (IL-2). Although IL-2 is indispensable for Treg, they cannot produce it themselves^7–9^. IL-2 levels in blood, cerebrospinal fluid, and inflamed tissues are typically far lower than the concentrations required to support Treg survival and function *in vitro*^10–15^. Moreover, circulating IL-2 has a very short half-life (6 to 20 minutes)^16^. Given this poor availability of IL-2, it is unclear how Treg access this critical resource *in vivo* at sites of autoimmunity.

Several cytokines are sequestered within the extracellular matrix (ECM) by binding to HS-containing proteoglycans (HSPG), including murine and human IL-2^17–21^. In addition, binding to HS has been shown to potentiate the impact of IL-2 on both antigen presentation^21^ and the pro-apoptotic effects of high dose IL-2^20^. However, many of these studies pre-date knowledge about FOXP3^+^ Treg. Unlike Treg, conventional T cells (Tconv) do not require exogenous IL-2 *in vivo*^22^, such that the functional relevance of tissue-bound IL-2 was unclear.

Here, we have tested the hypothesis that FOXP3^+^ Treg use heparanase (HPSE) to access HS-bound IL-2 from the ECM within inflamed tissues and that this supports Treg homeostasis and function. We have performed these studies in the experimental autoimmune encephalomyelitis (EAE) mouse model of multiple sclerosis, a setting where IL-2 and FOXP3^+^ Treg are known to modulate disease severity^23,24^. Our data reveal a previously unsuspected role for the tissue microenvironment in the regulation of Treg homeostasis and immune tolerance.

### IL-2 is bound to HS in EAE lesions

To explore the role of HS in autoimmune neuroinflammation, we asked whether IL-2 and HS are present at sites of inflammation in C57Bl/6/MOG_35-55_ -induced EAE. We first characterized the distribution of both IL-2 and HS in spinal cord tissue from naïve (healthy) and EAE animals. We observed that naïve spinal cord tissue has scattered IL-2 staining. This staining is not associated with HS staining, which is found mostly in vasculature and meninges (Fig. 1a). In contrast, spinal cord tissue from EAE mice shows increased IL-2 staining throughout areas of parenchymal inflammation, as well as increased HS deposition (Fig. 1a,b). Both IL-2 and HS staining were found on cellular structures with astroglial morphology (Extended Data Fig. 1, arrows), and in the form of diffuse staining in between cellular structures, suggesting association with peri- and extracellular matrix (Extended Data Fig. 1, asterisks). HS staining was also seen associated with some CD45 cells, both in perivascular spaces, as well as infiltrated into the CNS parenchyma. However, these cells generally did not stain positive for IL-2 (Extended Data Fig. 1, arrowheads). Staining for glial fibrillary acidic protein (GFAP), which stains the cytosekeleton of astrocytes, revealed that indeed, much of the cellular pattern of IL-2 and HS staining overlapped, or was directly adjacent to, GFAP+ structures, indicating that IL-2/HS deposition in EAE lesions is associated with reactive astrocytes (Extended Data Figs. 2 and 3).

**Figure 1.**
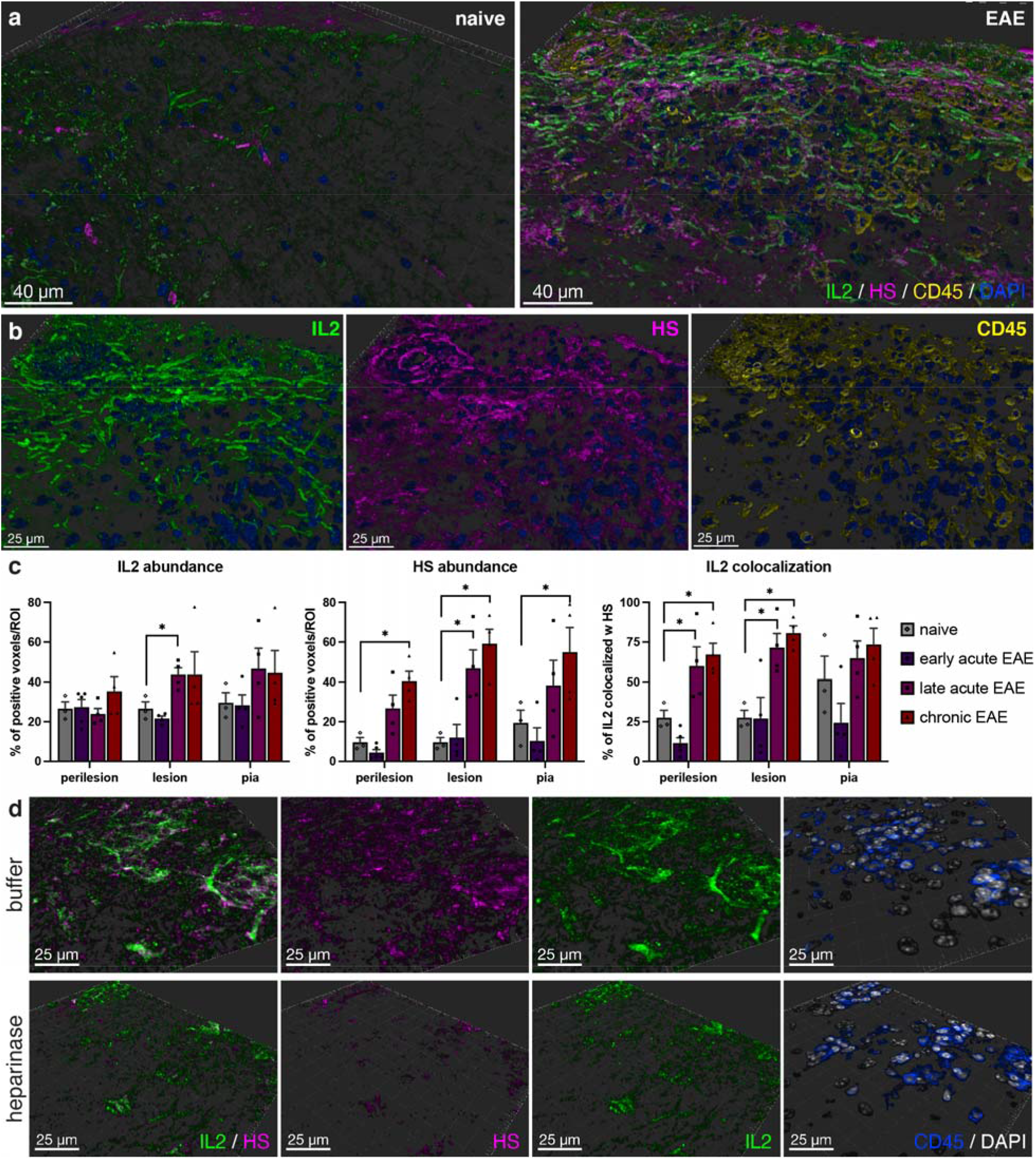
IL-2 and HS colocalize at sites of autoimmune neuroinflammation. **a**, Imaris surface rendering of immunofluorescent staining for IL-2 (green), HS (magenta) and CD45 (yellow) in naïve of EAE spinal cord tissue (29 days post immunization (dpi)). DAPI nuclear counterstain shown in blue. Representative of 3 separate EAE experiments. **b**, Detail of IL2 (green), HS (magenta) and CD45 (yellow) immunoreactivity in an EAE lesion. **c**, Imaris quantification of abundance of IL2 and HS immunoreactivity, and fraction of IL-2 colocalized with HS in naïve and EAE spinal cord tissue at different time points (early acute: 14 dpi, late acute: 29 dpi, chronic 40 dpi). Imaris colocalization analysis was done in areas surrounding CD45 cell infiltration (perilesion), areas with CD45 cell infiltration (lesion) and in the glia limitans underlying the meninges (pia). * P < 0.05, Mann-Whitney test, n = 3-5 separate areas analyzed per condition. **d**, Immunofluorescent staining for IL-2 (green) and HS (magenta) of a cerebellar EAE lesion that was treated with heparinase from *Flavobacterium heparinum* before immunofluorescent staining, or buffer treatment of a serial section as a control. CD45 (blue) staining is show to depict areas with immune cell infiltration. Representative of 2 separate experiments.

We then quantified the abundance and colocalization of IL-2 and HS throughout the course of EAE, using Imaris software. We observed that IL-2 staining was significantly increased in areas with CD45^+^ leukocyte infiltration (lesions), at the late acute stage of disease (Fig. 1c). HS was markedly increased in these areas as well, but also in areas adjacent to lesions (perilesion) and the glia limitans underlying the meninges, at the chronic stage of disease (Fig. 1c). This gradual increase correlated with the colocalization of IL-2 with HS, which was significantly increased in lesion, as well as perilesion areas at late acute and chronic stages (the percentage of Il-2 that is colocalized with HS ranging from 60 to 80%; Fig. 1c). These analyses show that both the abundance of HS and IL-2 increase over the disease course of EAE and that these co-localize extensively.

To determine whether the deposition of IL-2 deposition in inflammatory areas depends on its interaction with HS, we enzymatically removed HS structures using heparinase and subsequently assessed the presence of both IL-2 and HS in the treated tissue. We found that heparinase treatment not only removes HS from the tissue, but also reduces IL-2 immunostaining where it overlaps with HS (Fig. 1d), suggesting that IL-2 binds to HS in EAE lesions. Together, these data indicate that IL-2 is present and bound to HS at sites of autoimmune neuroinflammation.

### Binding to HS fragments increases IL-2 potency

We next asked whether binding to HS impacts the function of IL-2. We found that pre-incubation with soluble HS enhances the bioactivity of IL-2 as measured by CTLL2 proliferation (Fig. 2a). To specifically interrogate the role of the HS-binding capacity of IL-2 in this potentiating effect, we used a mutant of IL-2, T51P-IL-2, which contains a single point mutation in the HS binding domain of IL-2 that significantly decreases its ability to bind HS but leaves its soluble bioactivity intact^19^. Soluble IL-2 and soluble T51P-IL-2 have comparable effects on CTLL2 proliferation (Extended Data Fig. 4a). However, pre-incubation of IL-2 with soluble HS enhances its biological efficacy 4- to 100-fold, with only modest effects on T51P-IL-2 (Extended Data Fig. 4b). These data demonstrate that binding to soluble HS amplifies the bioactive potency of IL-2.

**Figure 2.**
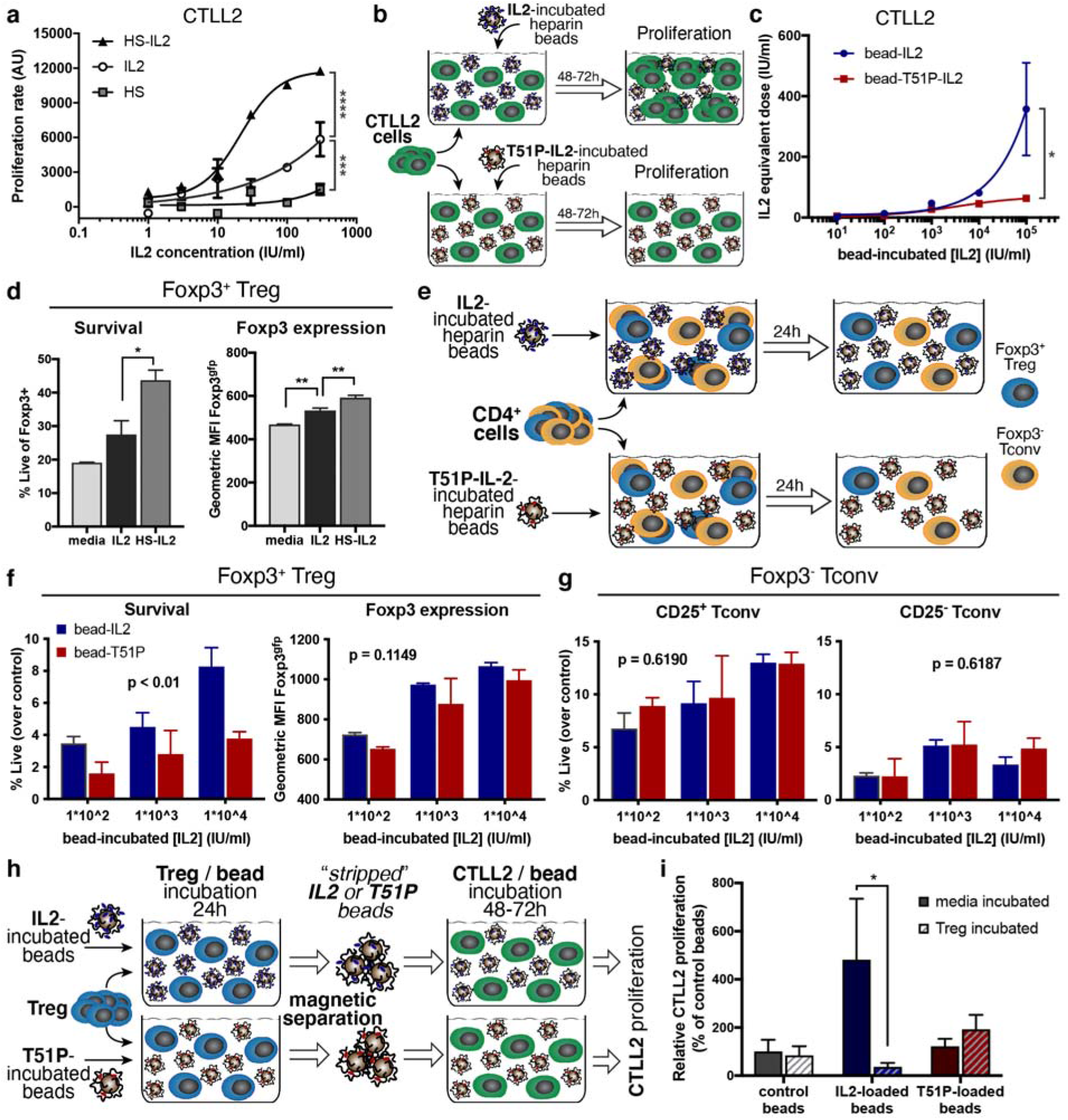
HS-bound IL-2 supports Treg homeostasis. **a**, CTLL2 proliferation, measured by resazurin reduction (arbitrary fluorescence units (AU)), in response to human recombinant IL-2 alone or pre-incubated with HS at a molecular ratio of 5:1. Equivalent doses of HS were used as a control. *** p < 0.001, **** p < 0.0001, two-way ANOVA. Shown is a representative of 4 independent experiments (mean +/- SEM of duplicate samples). **b**, Schematic overview of CTLL2 proliferation induced by heparin-coated beads pre-incubated with IL-2 or T51P-IL-2, an IL-2 mutant that has reduced binding to HS. **c**, CTLL2 proliferation in response to heparin-coated beads pre-incubated with IL-2 or T51P-IL-2. The concentration of IL-2 or T51P-IL-2 at which the beads were pre-incubated is depicted on the x-axis. Proliferation rate is shown as equivalent dose of soluble IL-2 or T51P-IL-2 at which a similar proliferative response is elicited as by the respective pre-incubated beads. Shown is a representative of 3 independent experiments (mean +/- SEM of triplicate samples). * p < 0.05, two-way ANOVA. **d**, Viability and FOXP3^gfp^ expression of induced FOXP3+ Treg, cultured in the presence of low-dose human recombinant IL-2 (20 IU/ml) alone, or IL-2 pre-incubated with HS, as described in Fig. 2A. Viability and FOXP3^gfp^ expression were measured by flow cytometry 72hr after start of induction. Shown is a representative of 5 independent experiments (mean + SEM of triplicate samples), * p < 0.05, ** p < 0.01, one-way ANOVA with Tukey’s multiple comparison correction. **e**, Schematic overview of the assay used to assess persistence of Treg induced by heparin-coated beads pre-incubated with IL-2 or mutant T51P-IL-2. **f and g**, Viability and FOXP3^gfp^ expression of FOXP3^+^ Treg (**f**) and (**g**) viability of CD25^+^ and CD25^-^ FOXP3^-^ Tconv, among CD4^+^ T cells cultured in the presence of heparin-coated beads pre-incubated with IL-2 or T51P-IL-2 (T51P). Cells were analyzed by flow cytometry 24 hr after start of culture. Percentage of viable cells among (**f**) FOXP3^+^ or (**g**) FOXP3^-^ cells is depicted, corrected for baseline viability of cells cultured in media only (without IL-2). All panels show representatives of 4 independent experiments (mean + SEM of triplicate samples); p-values depict variation due to the cytokine that the beads were incubated with (IL-2 vs. T51P-IL-2), determined by two-way ANOVA. **h**, Schematic overview of the assay designed to assess the effectiveness of Treg-mediated stripping of IL-2 from heparin-coated beads. Heparin-coated beads previously incubated with IL-2 or mutant T51P-IL-2 (T51P) were incubated with or without Treg overnight. These beads were then magnetically separated from Treg, washed, and added to CTLL2 cells. Proliferation of CTLL2 cells in response to residual bead-bound IL-2 was subsequently measured after 72 hours. **i**, CTLL2 proliferation in response to IL-2 and T51P-IL-2-pre-incubated beads cultured with or without Treg. Proliferation rate is shown as percentage of proliferation in response to control beads, that were incubated in media in both pre-incubation steps. Shown is a representative of 3 independent experiments (mean + SEM of 6 wells per sample). * p < 0.05, two-way ANOVA with Sidak’s multiple comparison correction.

To better assess the binding of IL-2 to HS and the biological implications of this binding, we coated magnetic beads with heparin, which is chemically identical to HS and only differs in the level of sulfation^25^. We then loaded these beads with increasing concentrations of IL-2 and T51P-IL-2 and washed off any unbound cytokine (Extended Data Fig. 5). Using CTLL2 cell proliferation as a read-out (Fig. 2b), we compared the efficacy of IL-2- and T51P-IL-2-loaded beads to that of soluble IL-2 and T51P-IL-2. We found that heparin-coated beads loaded with IL-2 deliver biologically active cytokine in a dose-dependent fashion, whereas the effective dose delivered by T51P-IL-2-loaded beads is much lower (Fig. 2a and Extended Data Fig. 4c).

### Treg utilize HS-bound IL-2 for homeostasis

Using FOXP3.GFP reporter mice, we next evaluated the impact of binding to HS on the ability of IL-2 to support *in vitro* homeostasis of CD4^+^FOXP3^+^ Treg. We first observed that pre-incubation of IL-2 with HS enhances its support of the survival and FOXP3 expression of *in vitro* induced Treg (Fig. 2d). Next, we used a competitive setting to compare the usage of HS-bound IL-2 between freshly isolated CD4^+^FOXP3^+^ Treg and CD4^+^FOXP3^-^ Tconv, using our heparin-coated beads (Fig. 2e). We found that beads incubated with IL-2, but not T51P-IL-2, support *in vitro* survival of FOXP3^+^ Treg in a dose dependent manner, although the effect on FOXP3 expression did not reach statistical significance (Fig 2f). We speculate that GFP protein may persist in cells as a “lagging indicator” of Treg homeostasis. In contrast, the viability of FOXP3^-^ Tconv was not affected by either IL-2 or T51P-IL-2-loaded heparin beads, regardless of whether they expressed the IL-2 receptor CD25^+^ or not (Fig. 2g).

To test whether Treg “strip” IL-2 from heparin-coated beads, we co-incubated Treg with heparin-coated beads that were pre-incubated with either IL-2 or T51P-IL-2. We then separated these beads from the Treg and subsequently co-incubated them with CTLL2 cells. By assessing the proliferative response induced by these beads we inferred how much IL-2 remained bound to these beads (Fig. 2h). As before, we found that IL-2-loaded beads, but not T51P-IL-2-loaded beads, induce proliferation of CTLL2 cells (Fig. 2i). However, beads that were previously co-incubated with Treg promoted far less proliferation, indicating that Treg are able to remove IL-2 bound to heparin. Together, these data indicate that Treg, but not Tconv, use exogenous, HS-bound IL-2 to support their homeostasis. Moreover, these data suggest that Treg actively “strip” IL-2 that is bound to sequestered HS.

### Treg express higher levels of HPSE than Tconv

The data above suggest that Treg may actively engage HS-containing structures to access bound IL-2. HPSE is the only known enzyme that specifically binds and cleaves HS chains^26^. It was previously shown that Tconv express HPSE and that it is bioactive^27,28^. However, a role for HPSE on Treg has not been studied. We therefore assessed the expression of HPSE in freshly isolated and sorted FOXP3^+^ Treg and FOXP^-^ Tconv, both in resting cells and after *in vitro* activation.

Using real-time RT-PCR we found that upon activation murine FOXP3^+^ Treg express markedly higher levels of *HPSE* than FOXP3^-^ Tconv (Fig. 3a). Western Blot analysis revealed a similar increase at the protein level (Fig. 3b,c). Moreover, addition of exogenous IL-2 to the culture media reduced the activation-induced upregulation of *HPSE* mRNA expression in Treg, suggesting that *HPSE* expression is regulated by the availability of IL-2 (Extended Data Fig. 6a).

**Figure 3.**
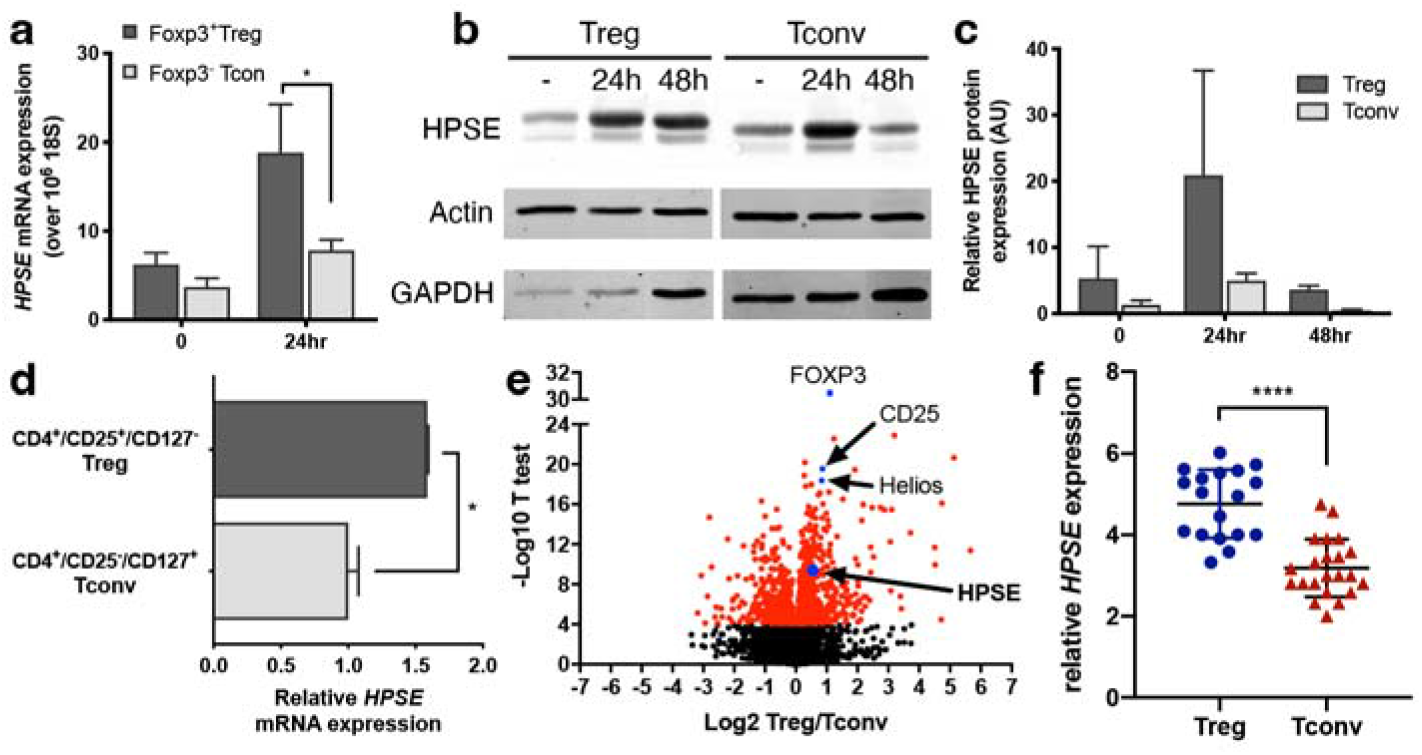
Enhanced HPSE expression after activation characterizes Treg. **a**, *HPSE* mRNA expression in FACS sorted murine FOXP3^+^ Treg and FOXP3^-^ Tconv after *in vitro* activation with aCD3/aCD28 (24h). Shown are mean relative expression + SEM of 3 independent experiments, compared to resting Tconv and normalized by *18S* mRNA expression. * p < 0.05, two-way ANOVA. **b**, Western blot analysis and **c**, semi-quantitation of HPSE protein expression in murine Treg and Tcon after *in vitro* activation with aCD3/aCD28. Actin was used as a control to normalize quantitation. **d**, *HPSE* mRNA expression in FACS sorted and *in vitro* activated human CD4^+^/CD25^+^/CD127^-^ Treg and CD4^+^/CD25^-^/CD127^+^ Tconv. Shown are mean relative *HPSE* expression +/- SEM of technical triplicates of a representative of 2 experiments, compared to Tconv and normalized by β*-actin* expression. * p < 0.05, unpaired two-tailed t-test. **e**, Volcano plot of statistical significance against fold change of genes differentially expressed between Treg and Teff isolated from human colon tissue. **f**, *HPSE* mRNA expression in FACS sorted CD4^+^/CD25^+^/CD127^-^ Treg and CD4^+^/CD25^-^/CD127^+^ Tconv isolated from human colon tissue. Shown are mean relative *HPSE* expression +/- SD, **** p < 0.0000001, unpaired two-tailed t-test.

We similarly observed higher expression levels of *HPSE* in human *in vitro* activated FACS-sorted CD4^+^/CD25^+^/CD127^-^ Treg, compared to sorted CD4^+^/CD25^-^/CD127^+^ Tconv (Extended Data Fig. 6b and Fig 3d). Moreover, RNA sequencing of FACS-sorted Treg and Tconv present in blood and colon tissue samples revealed that *HPSE* is differentially expressed in Treg compared to Tconv (Fig. 3e and Extended Data Fig 6d,e). Collectively, these data demonstrate that both Tconv and Treg upregulate *HPSE* upon activation but that only in Treg do HPSE levels remain elevated.

### HPSE enhances the ability of Treg to access HS-bound IL-2

We then sought to determine the functional impact of HPSE on the utilization of HS-bound IL-2. To this end, we first stably overexpressed human HPSE in CTLL2 cells, which already expressed low levels of murine HPSE (Fig. 4a). We then assessed their ability to respond to IL-2 bound to the heparin-coated beads described above. We found that HPSE overexpression results in an enhanced proliferative response to heparin-bound IL-2 compared to control CTLL2 cells (Fig. 4b).

**Figure 4.**
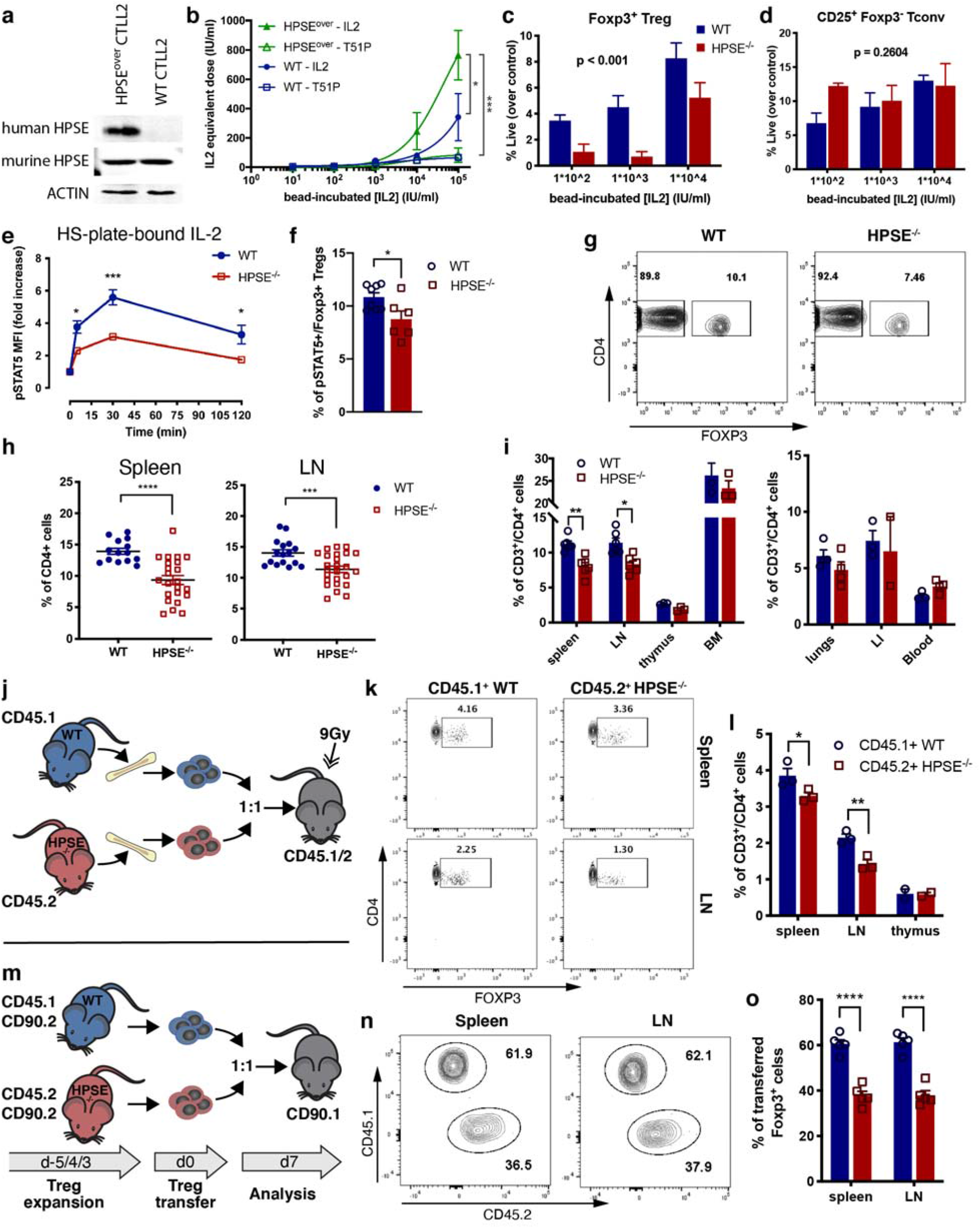
HPSE expression supports FOXP3+ Treg homeostasis *in vitro* and *in vivo*. **a**, Western blot analysis of human HPSE overexpression and mouse HPSE basal expression in CTLL2 cells transfected with HPSE construct (HPSE^over^), compared to untransfected CTLL2 cells (WT). **b**, Proliferation of wt CTLL2 cells (WT, blue) and HPSE-overexpressing CTLL2 cells (HPSE^over^, green) in response to heparin-coated beads that were pre-incubated with IL-2 (closed symbols) or T51P-IL-2 (T51P, open symbols). The concentration of IL-2 or T51P-IL-2 at which the beads were pre-incubated is depicted on the x-axis. Proliferation rate is depicted as equivalent dose of soluble IL2 or T51P-IL-2 at which a similar proliferative response is elicited as by the beads. Shown is a representative of 3 independent experiment (mean +/- SEM of triplicate samples). * p < 0.05, *** p < 0.001, variation due to the combination of the background of cells and the cytokine beads were pre-incubated with, determined with two-way ANOVA with Tukey’s multiple comparison correction. **c** and **d**, Viability of wt and HPSE^-/-^ FOXP3^+^ Treg (**c**) and CD25^+^ FOXP3^-^ Tconv (**d**) among CD4^+^ T cells cultured in the presence of heparin-coated beads pre-incubated with IL-2. Viability was measured by flow cytometry 24 hr after start of culture. Shown are representatives of 4 independent experiments (mean + SEM of triplicate samples); p-values depict variation due to the genotype (wt vs. HPSE^-/-^), determined by two-way ANOVA. **e**, STAT5 phosphorylation in wt and HPSE^-/-^ FOXP3^+^ Treg after stimulation with IL-2 that was sequestered by plate-bound HS. * p < 0.05, *** p<0.001, two-way ANOVA with Sidak’s multiple comparison correction. **f**, Percentage of pSTAT5+ cells among CD4^+^ FOXP3^+^ Treg isolated from naïve wt and HPSE^-/-^ spleen tissue. **g**, Representative flow cytometry plots depicting FOXP3^+^ Treg frequencies among CD4^+^ T cells in spleen tissue of wt and HPSE^-/-^ mice. **h**, Percentage of FOXP3+ Treg among CD4+ T cells in the spleens (lefts panel) and inguinal lymph nodes (right panel) of adult (3 to 6-month-old) wt and HPSE^-/-^ mice. Shown are mean + SEM, *** p < 0.001, **** p<0.0001, two-tailed t-test. **i**, Quantification of FOXP3^+^ Treg frequencies among CD4^+^ T cells in lymphoid (left panel) and non-lymphoid (right panel) tissues of wt and HPSE^-/-^ mice. BM, bone marrow; LI, large intestine. **j**, Schematic overview of competitive bone marrow transplantation of wt and HPSE^-/-^ donors. **k**, Representative flow cytometry plots depicting FOXP3^+^ Treg frequencies among transferred CD4^+^ T cells recovered from spleen and LN tissue of mice engrafted with bone marrow from wt and HPSE^-/-^ mice. **l**, Frequencies of FOXP3^+^ cells among wt and HPSE^-/-^ bone marrow-derived CD4^+^ T cells in lymphoid tissues of irradiated recipient mice. **m**, Schematic overview of competitive Treg transfer of wt and HPSE^-/-^ donors. **n**, Representative flow cytometry plots depicting CD45.1^+^ and CD45.2^+^ cell frequencies among transferred Treg recovered from spleen and LN tissue of recipient mice. **o**, Frequencies of CD45.1^+^ wt and CD45.2^+^ HPSE^-/-^ among Treg in lymphoid tissues of recipient mice. **g, i, l and o**, Shown are mean + SEM; * p < 0.05, ** p<0.01, two-tailed t-test with Holm-Sidak multiple comparison correction.

To interrogate the role of HPSE expression in Treg, we obtained HPSE deficient (HPSE^-/-^) mice and crossed these to the FOXP3.GFP reporter strain to generate HPSE^-/-^ -FOXP3.GFP mice. To facilitate comparisons of these strains, we backcrossed FOXP3.GFP.HPSE^-/-^ mice against FOXP3.GFP.HPSE^+/-^ mice for over 9 generations to generate matched FOXP3.GFP.HPSE^+/+^ controls. We then assessed the ability of freshly isolated Treg from these strains to access heparin-bead-bound IL-2 *in vitro*. We found Treg isolated from HPSE^-/-^ mice are unable to access bead-bound IL-2 as efficiently as Treg from WT mice (Fig. 4c). In contrast, neither CD25^+^ nor CD25^-^ Tconv are impacted by the loss of HPSE (Fig. 4d and Extended Data Fig. 7a), probably because they can produce IL-2 endogenously.

Given the importance of IL-2/STAT5 signaling in Treg homeostasis^29,30^, we next assessed the impact of HPSE expression on this signaling. We found that soluble IL-2 induces rapid STAT5 phosphorylation in both HPSE^-/-^ and WT Treg (Extended Data Fig. 7b). However, when IL-2 is delivered bound to HS-coated plates, STAT5 phosphorylation is significantly reduced in HPSE^-/-^ Treg, compared to WT Treg (Fig. 4e), indicating that Treg depend on HPSE to access and respond to HS-bound, but not soluble IL-2. Moreover, we found that the frequency of pSTAT5^+^ cells among Treg is reduced in spleen tissue from HPSE^-/-^ mice compared to WT mice (Extended Data Fig. 7c and Fig. 4f). This suggests that HPSE may promote tonic IL-2 responses *in vivo* as well.

### HPSE expression supports FOXP3^+^ Treg homeostasis in vivo

Next, we sought to define the role of HPSE in Treg homeostasis *in vivo*. We first examined the frequency of FOXP3^+^ Treg among CD4^+^ T cells in adult HPSE^-/-^ and WT mice. This revealed that Treg frequencies are significantly lower in secondary lymphoid tissue (spleen and lymph nodes) of HPSE^-/-^ mice, compared to WT mice (Fig. 4g,h). We observed the same pattern in both HPSE^-/-^.FOXP3.GFP on the C57Bl/6 background, as well as mice crossed with C57Bl/6 10BiT reporter mice from a different institution^48^ (Extended Data Fig. 7d). Consistent with this, FOXP3^+^ Treg frequencies are reduced in secondary lymphoid tissue of HPSE^-/-^ mice, but not in other tissues, including thymus, and the blood (Fig. 4i). Of note, the number and distribution of cytokine-producing CD4^+^ T helper cell subsets is not altered in lymphoid tissue of HPSE^-/-^ mice (Extended Data Fig. 7e), indicating that these effects are specific to Treg. Moreover, whereas the frequency of FOXP3^+^ Treg increases with age in wild type (WT) mice^31^, in HPSE^-/-^ mice, this aging-related increase in Treg frequency is significantly reduced (Extended Data Fig. 7f). Together, these data suggest that Treg homeostasis is impaired in the absence of HPSE.

To test whether the effect of HPSE expression on the homeostasis of Treg is cell intrinsic, we performed two competitive transfer experiments. First, we generated mixed bone marrow chimeras using congenic (CD45.1^+^) WT and (CD45.2^+^) HPSE^-/-^ donors and assessed the frequency of FOXP3^+^ Treg among donor-derived cells in lymphoid tissue of CD45.1/2 recipients (Fig. 4j). This revealed that Treg frequencies were significantly lower among CD4^+^ T cells derived from HPSE^-/-^ donor cells than those derived from WT donor cells in recipient secondary lymphoid tissue, but not the thymus (Fig. 4k,l).

Additionally, we FACS-sorted FOXP3^+^ Treg from CD45.1^+^ WT and CD45.2^+^ HPSE^-/-^ donor mice, both on the CD90.2 background, and adoptively transferred these cells into congenic CD90.1 recipient mice (Fig. 4m). Seven days post-transfer, the frequency of HPSE^-/-^ donor-derived cells among transferred FOXP3^+^ Treg was significantly lower than that of WT donor-derived cells in both spleen and lymph node tissue (Fig. 4n,o).

Together, these data indicate that the impairment seen in HPSE^-/-^ Treg homeostasis is intrinsic to hematopoietic cells. Moreover, our transfer studies with purified Treg establish that these effects have specific relevance to these cells in this model system.

### HPSE expression enables FOXP3^+^ Treg to access tissue-bound IL-2

Considering our observation that overlapping IL-2 and HS staining in EAE lesions appears to be astrocyte-associated (Extended Data Fig. 2 and 3), we next questioned whether Treg are capable of accessing IL-2 that is associated with astrocytes and whether HPSE is necessary for this. To test this, we pre-incubated confluent cultures of the astroglioma cell line U87-MG, which express HSPG on their cell surface^32^, with IL-2. After washing off any unbound IL-2, we co-cultured these with freshly isolated CD4^+^ T cells from WT and HPSE^-/-^ mice (Fig. 5a). We found that FOXP3^+^ Treg only survive in co-cultures with astrocytes when they have been pre-incubated with IL-2 and that this survival is significantly reduced in HPSE^-/-^ Treg compared to WT Treg (Fig. 5b). In contrast, the survival of CD25^+^ Tconv was much lower in these co-cultures and is not affected by HPSE expression (Fig. 5,c). Of note, the survival of CD25^-^ Tconv was supported by astrocyte cultures and is affected by HPSE expression to a small extent. However, this effect is not dependent on IL-2 (Extended Data Fig. 8a).

**Figure 5.**
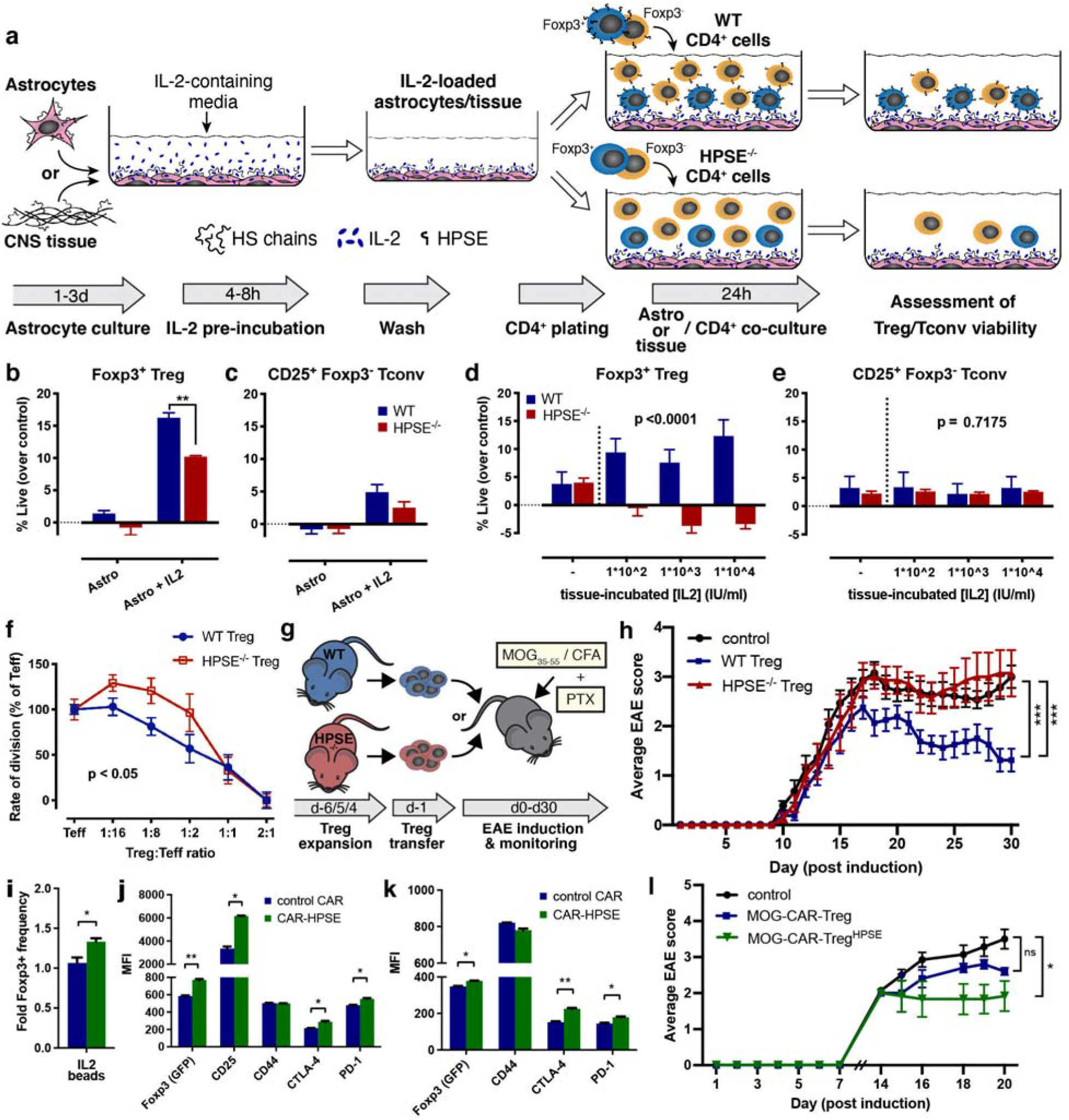
HPSE expression supports FOXP3+ Treg function *in vitro* and *in vivo*. **a**, Schematic overview of experiments designed to assess the ability of Treg to utilize IL-2 bound to U87-MG astrocytes or whole spinal cord (CNS) tissue in an HPSE-dependent manner. Astrocyte monolayers or spinal cord tissue (either freshly isolated and irradiated or snap frozen) from a wt mouse were pre-incubated with human recombinant IL-2 for 4-8 hr. After this cells/tissue were washed, removing any unbound IL-2, and subsequently co-incubated for 24 hours with CD4^+^ T cells isolated from wt or HPSE^-/-^ mice, after which viability of FOXP3^+^ Treg and FOXP3^-^ Tconv was assessed by flow cytometry. **b** and **c**, Viability of wt and HPSE^-/-^ (**b**) FOXP3^+^ Treg and (**c**) CD25^+^ FOXP3^-^ Tconv among CD4^+^ T cells co-cultured with U87-MG cells that were pre-incubated with IL-2 or media alone as a control. **d** and **e**, Viability of wt and HPSE^-/-^ (**d**) FOXP3^+^ Treg and (**e**) CD25^+^ FOXP3^-^ Tconv among CD4^+^ T cells co-cultured with mouse spinal cord tissue that was pre-incubated with IL-2 or media alone as a control. (**b-e**) Percent of viable cells among FOXP3^+^ or FOXP3^-^ cells is depicted, corrected for baseline viability of cell cultured in media alone. All panels show mean + SEM of triplicate samples of a representative of 4 (**b** and **C**) or 3 (**d** and **e**) independent experiments. (**b** and **c**) * p < 0.05, ** p<0.01, two-tailed t-test with Holm-Sidak multiple comparison correction. (**d** and **e**) p-values depict variation due to genotype (wt vs. HPSE^-/-^), determined by two-way ANOVA. **f**, *In vitro* suppression of wt effector (FOXP3^-^) T cells (Teff) by wt and HPSE^-/-^ FOXP3^+^ Treg. The rate of division of Teff is plotted as percentage of unsuppressed Teff. Shown are mean +/- SEM of triplicate wells of a representative of 2 independent experiments. P-value shown: two-way ANOVA between wt and HPSE^-/-^ Treg. **g**, Schematic overview of transfer of *in vivo* expanded, CD4^+^/CD25^+^ sorted, Treg into wt recipients and subsequent EAE induction. **h**, *In vivo* suppression of EAE by wt and HPSE^-/-^ FOXP3^+^ Treg that were transferred 1 day prior to induction of EAE. Shown is average disease severity +/- SEM of all animals, **\***\* p < 0.01, **** p<0.0001, Friedman test. Shown is a representative of 2 independent experiments. **i**, Frequency and (**j**) expression levels of Treg functional markers of FOXP3+ cells among total CD4^+^ cells transfected with HPSE-CAR after 72h culture with IL-2-incubated heparin beads. Shown are mean + SEM (triplicate samples) of the fold change of percentage FOXP3^+^ cells, relative to control CAR cultured without IL-2, and of geometric MFI of each marker. **k**, Expression levels of Treg functional markers on FOXP3+ cells recovered 3 days after adoptive transfer into CD45.2 recipients (*n* = 3-4 animals per group). **i**, * p < 0.05, two-tailed t-test (**j** and **k**) ** p<0.01, **** p<0.0001, two-way ANOVA with Sidak’s multiple comparison correction. **l**, *In vivo* suppression of EAE by control and HPSE-overexpressing MOG-CAR Treg that were transferred on day X after induction of EAE (*n* = 5-6 animals per group). Shown is average disease severity +/- SEM of all animals, **\*** p < 0.05, Friedman test.

We also tested whether Treg are capable of utilizing IL-2 associated with CNS tissue. To this end, we pre-incubated homogenized spinal cord tissue from healthy mice with IL-2 and, after washing off unbound IL-2, co-cultured this with CD4^+^ T cells. Using these preparations, we observed that IL-2 pre-incubated spinal cord tissue supports the survival of WT Treg in an IL-2 dose-dependent manner and that this is significantly reduced in HPSE^-/-^ Treg (Fig. 5d). Neither CD25^+^ nor CD25^-^ Tconv survival is enhanced by IL-2 pre-incubation of spinal cord tissue and does not depend on HPSE (Fig. 5e and Extended Data Fig. 8b). Consistent with this, HPSE-overexpressing CTLL2 cells are also able to proliferate, in a dose-dependent manner, in response to spinal cord tissue pre-incubated with IL-2, but not in response to T51P-IL-2 pre-incubated spinal cord tissue (Extended Data Fig. 8c,d). However, control CTLL2 do not display this proliferative response to tissue-associated IL-2 (Extended Data Fig. 8c,d), suggesting that perhaps, in contrast to the assays with heparin-beads or HS-incubated IL-2 described before, expression of HPSE is critical for CTLL2 cells to access IL-2 that is bound to tissue.

Together, these data indicate that Treg are capable of accessing IL-2 that is associated with cells or tissue and that this is dependent on HPSE expression.

### HPSE expression supports FOXP3^+^ Treg function in vitro and in vivo

Finally, we questioned whether HPSE expression also affects Treg function in suppressing immune responses. We found that HPSE^-/-^ Treg have impaired suppressive function relative to WT Treg *in vitro* (Fig. 5f). Furthermore, we observed that adoptively transferred HPSE^-/-^ Treg are not able to suppress autoimmune neuroinflammation *in vivo* in the EAE mouse model of MS, whereas WT Treg are (Fig. 5g,h, Extended Data Fig. 8e,f and Table 1). This reduced disease severity is mainly due to a higher recovery rate (more animals converting to a lower disease score) in mice that received WT Treg than in mice that either received HPSE^-/-^ or no Treg (Extended Data Fig. 8f), consistent with the notion that Treg are involved in the recovery of EAE^33,34^.This treatment effect was correlated with a change in the distribution of immune cells infiltrating the spinal cord of these mice, as assessed by flow cytometry (Extended Data Fig. 8g,h). Spinal cords of animals that received WT Treg contained markedly fewer infiltrating myeloid cells, which are the main drivers of disease progression in EAE^35,36^. This suggests that the therapeutic effect of transferred WT Treg extends to generally reducing inflammation and supporting greater recovery.

Together, these data indicate that HPSE facilitates the suppressive function of Treg *in vitro* and *in vivo*.

### HPSE over-expression supports the ability of FOXP3^+^ Treg to access HS-bound IL-2

Finally, to explore the translational potential of our findings, we assessed the impact of HPSE overexpression on the stability and function of chimeric antigen receptors (CAR) Treg stability and function, which show promise for the treatment of autoimmune diseases, such as MS^37,38^. For this, we used the flexible mAb-directed CAR clone 1X9Q, which contains an anti-FITC scFv portion fused to CD28 and CD3? costimulatory domains^39^, and fused the expression sequence of HPSE to this construct, as a proof-of-concept (Extended Data Fig. 9a). We first transfected mRNA produced from these constructs into CD4^+^ T cells from WT C57Bl/6 mice, resulting in high transfection efficiency for both constructs (Extended Data Fig. 9b). When cultured with IL-2-loaded heparin-beads, we observed that persistence of FOXP3^+^ cells is marginally, but significantly, higher among these HPSE overexpressing CAR T cells than control CAR T cells (Extended Data Fig. 9c and Fig 6i). Moreover, cultured HPSE overexpressing CAR Treg have higher expression of not only FOXP3, but also the functional Treg markers CTLA-4 and PD-1 (Fig. 6j). Similarly, when CD45.1 FOXP3^+^ HPSE overexpressing CAR Treg were adoptively transferred into CD45.2 mice and recovered 3 days later, they exhibited slightly increased FOXP3 and co-inhibitory marker expression (Fig. 6l and Extended Data Fig. 9d,e). Finally, we targeted HPSE overexpressing CAR Treg towards myelin by incubating them with a FITC-conjugated monoclonal antibody against MOG, before transferring them into EAE animals and testing their ability to suppress disease. Transferring these MOG-specific CAR Treg into mice at peak EAE revealed that overexpression of HPSE increased their ability to reduce disease severity compared to control CAR Treg (Extended Data Fig. 9f and Fig. 6k). Together, these data suggest that the expression of HPSE enhances the phenotypic stability and function of CAR Treg.

## Discussion

In this study, we have examined how FOXP3^+^ Treg obtain IL-2 at sites of autoimmune neuroinflammation. We report that IL-2 is sequestered by HS within the ECM and that FOXP3^+^ Treg use HPSE to access this essential resource to sustain their survival and function *in vivo*. In this way, ECM-bound IL-2, sequestered within tissues at sites of inflammation, may sustain regulatory responses in post-inflammatory settings. This mechanism may be particularly important in preventing and/or resolving autoimmunity, where Treg survival and function are critical for the maintenance of immune homeostasis.

Our finding suggest that tissue interactions are vital to the effects of IL-2 on Treg. Binding to soluble HS fragments, which could be generated locally *in vivo* due to cleavage of ECM HS chains by HPSE, enhances the biological effect of IL-2 on both FOXP3^+^ Treg and CTLL2 cells, a T cell line exquisitely dependent on exogenous IL-2. Consistent with these effects, HS was reported to potentiate the effects of IL-2 on Tconv proliferation^20^. HS similarly potentiates the impact of other cytokines; HS-binding of IL-12 significantly reduces the EC_50_ of this cytokine by acting as a co-receptor which enhances interactions between IL-12 and its receptor subunits^40^. In light of this potentiating effect of HS, and given the low levels of soluble IL-2 levels in blood, cerebrospinal fluid, and other fluids^10–14^, HS-bound IL-2, liberated by HPSE, may provide a critical source of support for Treg homeostasis *in vivo*. To be clear, we do not claim that all IL-2 is associated with HS. Rather, we propose that by influencing the spatial and temporal properties of IL-2 in tissues, the ECM is a partner in local immune homeostasis. These findings have clear implications for IL-2 as a therapeutic in autoimmunity and other inflammatory settings, and may inform future endeavors to design synthetic IL-2 or low-dose IL-2 treatment protocols, which have shown much promise in clinical trials^41–43^.

Competition for IL-2 is one of the mechanisms by which Treg suppress T-cell responses^29,44^. Therefore, the differential HPSE expression between Treg and Tconv may provide Treg with a competitive edge over effector T cells at sites of inflammation, where much of IL-2 is sequestered to HS in the ECM. Consistent with this, HPSE expression does not appear to be a benefit to Tconv, but rather play a critical role in Treg homeostasis and function.

The source of IL-2 in the CNS in the EAE model may be astrocytes given the colocalization of IL-2 with GFAP+ astrocytes in lesions. Consistent with this, astrocytes are reported to be able to produce both IL-2 and HS^32,45–47^. However, T cells, neurons and other cell types can also produce IL-2 and make contribute to the presence of IL-2 in these environments.

In this work, we have focused our efforts on targeted Treg studies, because of the complex phenotypes associated with systemic loss or inhibition of HPSE. However, HPSE is expressed by other cell types, including innate and adaptive immune cells^26,48,49^ and may impact these cells in distinct ways. Future studies will interrogate these connections. Likewise, distinct effects of HPSE on different cell types may underlie the conflicting data on HPSE inhibitors, some of which also have bioactivity independent of effects on HPSE^50^. Several such small molecules are promising therapies in cancer^51^ while at least one was shown to worsen EAE^52^. It has also been shown that administration of exogenous HPSE reduces disease in EAE and increases the production of anti-inflammatory cytokines by T cells^53^. Overall, studies on the role of HPSE in MS and EAE are limited and challenging to interpret, given the multiple cell types and tissues involved. Further in-depth studies involving cell- and tissue-specific approaches are warranted to fully uncover the role of HPSE in MS.

Building on the insights described here, we provide proof-of-principle data that (CAR) Treg engineered to express HPSE are better able to utilize HS-bound IL-2 and have enhanced phenotypic stability *in vitro* and *in vivo*. MOG-specific CAR Tregs have previously been shown to diminish disease severity in EAE upon intranasal transfer^54^. The results we present here demonstrate the promise of considering the cell-ECM interphase in attempts to enhance the function of (CAR) Treg based therapies.

To conclude, we demonstrate that IL-2 is sequestered by HS at sites of autoimmune inflammation, that HS-bound IL-2 promotes FOXP3^+^ Treg survival and function, and that Treg require HPSE to access this essential resource. These interactions resolve a critical gap in our understanding of Treg homeostasis and function *in vivo*. Therefore, these insights may provide new avenues for improving IL-2 and/or Treg treatment protocols.

## Acknowledgments

We thank Dr. Manish Butte for critical reading of the manuscript and the Stanford Shared FACS Facility for technical assistance.

## Funding

This work was supported by

National Institutes of Health grant R01DK046635 (GTN)

National Institutes of Health grant K08DK080178-05 (PLB)

National Institutes of Health grant R01DK096087-01 (PLB)

National Institutes of Health grant U01AI101984 (PLB)

National Institutes of Health grant R21AI133240-01A1 (PBL, HFK)

National MS Society Pilot Research Grant PP-1503-03972 (PBL, HFK)

CSGADP Pilot Award under NIH grant UO1AI101990 (TNW)

BIRT supplement AR037296 (TNW).

Stanford University Institute for Immunity, Transplantation and Infection Young Investigator Award (HFK)

Juvenile Diabetes Research Foundation grant nPOD 25-2010-648 (TNW)

The Center for Translational Research at BRI (GTN)

Helmsley Charitable Trust nPOD Award for Team Science (TNW)

## Author contributions

Conceptualization: HFK, IK, PLB

Methodology: HFK, IK, GK, BAF

Investigation: HFK, HAM, IK, GK, BAF, JK., COM, KRB, MGK, MP-C, S-WT

Resources: IV, J-PL, DP, EHM, DAH, KCG, TD LS

Writing – Original Draft: HFK

Writing – Review & Editing: HFK, HAM, IK, PLB

Supervision: GTP, TNW, PLB

Funding Acquisition: HFK, GTN, TNW, PLB

## Extended Data

Materials and Methods

Extended Data Figures 1-9

Extended Data Table 1

