## Supplementary material for "FOXP3^+^ regulatory T cells use heparanase to access IL-2 bound to ECM in inflamed tissues": Methods, Extended Data 1-13

**This PDF file includes:**

Materials and Methods
Extended Data Figures 1 to 13
Extended Data Table 1
Extended Data References

### Materials and Methods

**Mice.** To generate HPSE$^{-/-}$-FOXP3.GFP mice, harboring a GFP-FOXP3 fusion reporter knock-in allele in addition to a targeted disruption of the HPSE gene, HPSE$^{-/-}$ mice, described before[1] were crossed with FOXP3.GFP reporter mice[2], a kind gift of Dr. Alexander Rudensky. Both strains were maintained on the C57BL/6J background and offspring was backcrossed and maintained as a homozygous line. In addition, resulting HPSE$^{-/-}$-FOXP3.GFP mice were crossed with 10BiT-FOXP3.GFP mice, derived from crossing 10BiT mice containing a Thy1.1 (CD90.1) reporter under the control of the IL-10 promoter and FOXP3.GFP reporter mice[3], a kind gift of Dr. Casey Weaver. Wild-type (CD45.2) and congenic CD45.1 C57BL/6J, C.129P2(B6)-Il2$^{tm1Hor}$/J (IL2$^{-/-}$) and B6.129S7-Rag1$^{tm1Mom}$/J (RAG1$^{-/-}$) mice were obtained from Jackson Laboratories (Bar Harbor, ME). Congenic CD90.1 C57BL/6J were a kind gift of Dr. Robert Negrin. All mice were maintained in specific pathogen-free AAALAC-accredited animal facilities at the BRI and Stanford University and handled in accordance with institutional guidelines.

**Induction of EAE.** EAE was induced in female or male C57BL/6J mice (Jackson Laboratories) at 8-12 weeks of age by subcutaneous immunization with an emulsion containing 200 µg of myelin oligodendrocyte glycoprotein (35–55) ($MOG_{35–55}$) in saline and an equal volume of complete Freund's adjuvant (CFA) containing 400 ng of mycobacterium tuberculosis H37RA (Difco Laboratories, Detroit, MI). All mice were administered 400 ng of pertussis toxin (List Biological, Campbell, CA) intraperitoneally (i.p.) at 0 and 48 h post-immunization. Mice were weighed and monitored daily for clinical symptoms and scored as follows: 0, no clinical disease; 1, tail weakness; 2, hindlimb weakness; 3, complete hindlimb paralysis; 4, hindlimb paralysis and some forelimb weakness; 5, moribund or dead. At peak of disease (day 22), mice were sacrificed and brain tissue was harvested for immunohistochemical analysis.

**Histology.** Mice were deeply anesthetized with ketamine/xylazine (100 mg/kg and 7 mg/kg, respectively), transcardially perfused (saline followed by 4% paraformaldehyde in PBS) and spleens, lymph nodes (inguinal, axillary and brachial) and brains were collected. Tissue was post-fixed overnight in 4% paraformaldehyde at 4 °C, then cryoprotected in 30% sucrose and stored at 4 °C until they were embedded in Optimal Cutting Temperature compound (Tissue-

Tek) and frozen at −80 °C. Cryosections (20 µm) were stained using a free-floating immunohistochemistry protocol. Briefly, after blocking in 5% normal donkey serum (NDS) in Tris-buffered saline (TBS) with 0.3% Triton X-100 (TBS-T), sections were incubated overnight at 4 °C in TBS-T with 1% NDS and the following primary antibodies: mouse anti-HS (clone F58-10E4, Seikagaku; 1:50), chicken anti-IL2 (Sigma GW22461F; 1:250) and rat anti-CD45 (clone 30-F11, Life Technologies MCD4500; 1:500). After 3 washes with TBS-T, sections were incubated with species-specific fluorescent secondary antibodies from donkey conjugated with Alexa Fluor 488, 594, or 647 (Jackson Laboratories; all 1:500) for 45 min at RT.

**Imaging and image processing.** All images were acquired using an automated Zeiss CellDiscover7 LSM900 confocal microscope (Zeiss), using identical settings across all samples. z-stacks were acquired of regions of interest (inflammatory lesions or corresponding areas in control tissue) using an 20×/0.95 NA objective, at a voxel size of 0.097 x 0.097 x 0.63 µm. Channels were acquired sequentially to prevent spill-over. Images we pre-processed in Fiji (version 2.3.0; National Institutes of Health) using rolling ball background subtraction, with a rolling ball radius of 50.0 pixels. Preprocessed images were then analyzed using Imaris software (Bitplane). First, masks were made for each channel to enable surface rending and colocalization analysis, using identical thresholding and voxel minimum parameters for all samples. In addition, regions of interest (ROIs) were set across multiple z planes for areas with CD45 cell infiltration ("lesion"), areas adjacent to lesions ("peri-lesion") and the glia limitans underlying the meninges ("pia"). Then colocalization of IL-2 and HS fluorescence was calculated using the Imaris colocalization algorithm within these ROIs. Data is represented as % of positive IL-2 or HS voxels within the ROI, as well as % of IL-2 positive voxels that are colocalized with HS.

**Heparinase treatment of tissue.** 20 µm cryosections were incubated with 50 mU Heparinase III from *Flavobacterium heparinum* (Sigma-Aldrich) in 10 mM HEPES and 2mM CaCl2, pH7.0 at 37C for 1 hr. As a control, tissue was incubated at the same conditions in buffer without heparinase. After HS digestion, sections were washed once in PBS and stained as described above.

**CTLL2 culture and resazurin proliferation assays.** For maintenance, CTLL2 cells were cultured in media (RPMI 1640 containing 2.05 mM L-Glutamine, 10% FBS and 1x penicillin/streptomycin; GE Healthcare, Pittsburgh, PA) supplemented with recombinant human IL-2, produced in HEK cells, at EC75, which was established measuring proliferation in a dose-response assay. For proliferation assays, CTLL2 cells were first washed twice with media not containing IL-2 and incubated in media not containing IL-2 for at least 4 hours at 37˚C. Cells were then washed again and plated at a density of $0.5x10^5$ cells per well in a total of 100 µl media. After 24 to 48 hours, proliferation was assessed by reduction of resazurin to resorufin[4]. Resazurin (Acros Organics) stock solutions were made in RPMI 1640 at 5mg/ml (w/v), filter sterilized and stored at -20˚C until use. Resazurin was added at a final concentration of 1 mg/ml to the cell cultures for the last 8 to 16 hours of cell culture. Fluorescence of the resorufin was then measured at 560 excitation / 590 emission using a Spark multimode microplate reader (Tecan).

**CTLL2 - tissue co-culture.** Spleen and lymph node tissue was harvest from naïve C57Bl/6 or $RAG1^{-/-}$ mice, or mice immunized by subcutaneous injection of a similar CFA preparation as used for induction of EAE. The tissue was then either snap frozen first and processed upon thawing, or processed immediately and frozen afterwards. Tissue was mechanically dissociated in RPMI 1640 media using a dounce homogenizer. IL-2 starved CTLL2 cells (culture for 4-8hrs without IL-2) were then co-cultured with tissue lysates and proliferation of CTLL2 cells was analyzed as described above. Where indicated, tissue was digested with 50 mU Heparinase I, II and III from *Flavobacterium heparinum* for 1 hr at 37˚C and subsequently washed before coculture, and the IL-2 receptor antagonist H9-RETR was added at 1 µM to co-cultures.

**Generation of IL-2 and T51P-IL-2.** Human recombinant IL2 and the IL-2 analog in which the threonine at position 51 was replaced by a proline (T51P-IL-2) were expressed using a baculovirus system in Hi5 insect cells and purified as previously described[5]. Full vector sequences are available upon request.

**HS preincubation of IL-2.** Human recombinant IL-2 (either Proleukin (Novartis Pharma), or in-house generated IL-2) was co-incubated at a molecular ratio of 5:1 with heparan sulfate (Sigma Aldrich) for 8 hrs at RT, in a minimal volume of PBS. Aliquots were frozen at -80˚C and diluted to working concentration in media prior to experiments.

**Generation and IL-2 loading of heparin-coated magnetic beads.** Thiolated heparin (BioTime Inc.) was coupled to 1 μm BcMag™ epoxy-activated magnetic beads (Bioclone Inc.) as per manufacturer's directions to yield heparin-coated beads. These beads were then incubated with IL-2 or T51P-IL-2 in cell culture media (RPMI 1640 containing 10% FCS and penicillin/streptomycin) at indicated concentrations. Beads were washed 3 times with cell culture media before co-culture with cells at a 1:5 cells:beads ratio.

**Isolation and culture of mouse CD4$^+$ T cells and Treg.** Total leukocytes were isolated from inguinal, axial, brachial and mesenteric lymph nodes and spleen cells from 8 to 12-week-old mice. Tissue was homogenized through a 70 μm strainer and red blood cells were lysed in the splenocyte suspensions. CD4+ T cells were isolated from the pooled cell suspensions using an EasySep™ Mouse CD4+ T Cell Isolation Kit (Stemcell Technologies, Vancouver, Canada) following the manufacturer's instructions. Treg were isolated from total cell suspensions using an EasySep™ Mouse CD25 Regulatory T Cell Positive Selection Kit (Stemcell Technologies). Cells were cultured in T cell media (RPMI 1640 containing 2.05 mM L-Glutamine, 10% FBS, 1x penicillin/streptomycin, 50μM β-mercaptoethanol, and 1mM sodium pyruvate), supplemented with IL-2 when indicated. For real-time PCR analysis of HPSE expression, FOXP3$^{GFP+}$ Treg were sorted from CD4+ T cell preparations using a BD FACSAriaII cell sorter. Viability analysis of CD4$^+$ T cells under the experimental conditioned described in the figure legends was performed by staining with the amine-reactive dyes Zombie Red or Zombie Aqua (Biolegend), according to manufacturer's instructions, and flow cytometry.

**Flow cytometry.** Single cell suspensions were prepared from spleen, lymph nodes, thymus and femurs of mice. Lungs and large intestines were minced and digested with 100IU/ml Collagenase I and 25 IU/ml DNase I (both from Roche). Red blood cells were lysed in red blood cell lysis buffer (Sigma-Aldrich). For phosphoflow staining, cells were fixed by resuspending in 50 μl pre-warmed of Cytofix Fixation Buffer (BD Biosciences) and incubated at 37°C in a $CO_2$ incubator for 15 minutes. Cells were then washed in FACS buffer (3% FBS (w/v) and 2 mM EDTA in PBS) and cell surface antigens were stained with anti-CD3-BV785 (17A2), anti-CD4-BV421 (RM4-5) and anti-CD25-APC-Cy7 (PC61) in FACS buffer for 30 min on ice. After washing, cells were then resuspended in pre-chilled Phosflow Perm Buffer III (BD Biosciences) and incubated on ice for 30 minutes. The cells were washed two more times with FACS buffer prior to staining using an antibody against pSTAT5 (Y694, BD Biosciences), diluted in FACS stain buffer on ice for 60 minutes. After staining, cells were washed once, resuspended in FACS stain buffer and analyzed. For cell viability analysis, $1x10^6$ cells were stained with anti-CD3-BV785 (17A2), anti-CD4-BV421 (RM4-5), anti-CD25-APC-Cy7 (PC61; all from BioLegend) and ZombieRed Live/Dead dye (Biolegend). For phenotyping and cytokine staining $5x10^6$ cells were stimulated with 50ng/ml PMA and 500ng ionomycin (Sigma-Aldrich) in the presence of Brefeldin A (Biolegend) for 5 hours. Cells were then stained with anti-CD3-PE-Cy7 (17A2), anti-CD4-BV785 (RM4-5), and anti-CD8-BV711 (53-6.7), fixed using IC buffer (Thermo Fisher Scientific), permeabilized with 0.5% saponin in PBS (w/v) and stained with anti-IFNγ-APC (XMG1.12), anti-IL-17-PE (TC11-18H10.1) and anti-TNFα-PE (MP6-XT22). Flow cytometry was performed on an LSRII (Becton Dickinson) in the Stanford Shared FACS Facility and data analysis (including the built-in tSNE plugin )was done using FlowJo 9 or 10 software (Treestar).

**Cell sorting.** For mouse T cell sorting, cell suspensions were made from spleen and lymph node tissue. For human T cell sorting, PBMC were prepared from whole blood using Percoll gradient (GE Healthcare). Mouse and human $CD4^+$ cells were enriched using an EasySep Mouse or Human CD4 isolation kit (Stemcell Technologies). Mouse cells were then sorted based on FOXP3.GFP expression. Human cells were stained with anti-CD4-FITC (RPA-T4; Biolegend), anti-CD127-PerCpCy5.5 (A019D5) and anti CD25-PE-Cy7 (M-A251, both BD Pharmingen) prior to sorting. Cells were sorted using a FACS Aria III (BD Biosciences) in the Stanford Shared FACS Facility and lysed in Trizol (Thermo Fisher Scientific) for RNA expression analysis.

**In vitro activation of T cells.** 96 well flat bottom tissue culture plates were pre-coated with 2.5 µg/ml anti-CD3 Ab (clone 145-2C11; Biolegend, San Diego, CA). FACS-sorted mouse $CD4^+/FOXP3^+$ or $CD4^+/FOXP3^-$ cells, and human $CD4^+CD127^-CD25^+$ Treg or $CD4^+CD127^+CD25^-$ Tconv (for RNA analysis), or bead-sorted mouse $CD25^+$ or $CD25^-/CD4^+$ cells (for protein analysis) were plated at a density of $2.5x10^5$ per well in T cell media containing 0.5 µg/ml anti-CD28 (clone 37.51; Biolegend). No exogenous IL-2 was added unless otherwise noted.

**Real-time PCR**. After *in vitro* activation, cells were harvested and RNA was collected using an RNeasy mini kit (Qiagen). cDNA was prepared from 350ng total RNA reverse transcribed in a 40ul reaction mix with random primers using the High-Capacity cDNA Reverse Transcription Kit (Applied Biosystems), according to manufacturer's instructions. 1.2ul cDNA was amplified in 1XTaqman Fast Universal PCR Mix with 250nM Taqman probe (all Applied Biosystems) in a 20ul reaction using the Fast program for 50 cycles on an ABI7900HT thermocycler. All samples were run in duplicate and data were analyzed using the Comparative Ct Method with software from Applied Biosystems. Estimated copy numbers were generated from a standard curve created by using a selected reference cDNA template and Taqman probe. For human T cells, qPCR was performed using SYBR® Green PCR Master Mix (Applied Biosystems). Primer sequences are available upon request.

**Western Blot.** After *in vitro* activation, $CD25^+$ or $CD25^-/CD4^+$ cells were harvested and taken up in RIPA buffer with HALT protease and phosphatase inhibitor mix (Thermo Fisher Scientific). Lysates were then incubated on ice for 30 minutes, spun down at 13,000g for 30 min at 4˚C and supernatants were transferred to a new eppendorf tube. Protein content was measured using a BCA assay (Pierce), according to manufacturer's directions. 25 ug of total protein was loaded onto a 12% PAGE gel. Proteins were transferred to nitrocellulose membranes and immunoblotted with primary antibodies against murine HPSE (rabbit antibody PAA711Mu04, CloudClone Corp.), human HPSE (mouse antibody clone HP3/17, ProSpec) and murine β-actin (rabbit antibody Poly6221, Biolegend) and subsequently DyLight-680 or DyLight800-conjugated secondary antibodies. Antibody binding was visualized and quantified using and Odyssey Imaging system and Image Studio software (Li-Cor).

**RNA sequencing of human donor derived T cells.** Approval for blood and colon tissue collection was obtained through the Benaroya Research Institute Institutional Review Board (No. 10090). All patients signed written informed consent documentation prior to collection of samples. Surgically resected tissue was promptly transported to the laboratory in cold phosphate-buffered saline and washed of blood, mucous, and stool in HBSS. Colonic mucosa was sharply dissected away from underlying fat and muscle and subjected to 3 washes of HBSS containing EDTA and DTT to remove mucous and epithelium. Residual lamina propria (LP) was then rinsed with RPMI, chopped into fine pieces, and digested for 90 minutes with collagenase and

DNAse in RPMI supplemented with MgCl2 and CaCl2. The fluid phase of this digest was then filtered through a 100-mm screen, centrifuged, and washed in phosphate-buffered saline. LP cells were frozen in 7% DMSO in fetal calf serum in a Mr. Frosty freezer jar placed at -80°C and then transferred to liquid nitrogen. Homogenized LP cells were later thawed and stained with viability dye (ViaProbe) and antibodies to HLA-DR (BioLegend, clone L243), CD11c (eBioscience, clone Bu15), CD4 (BDBioscience, clone RPA-T4), CD45RA (BD Bioscience, clone HI100), CD19 (BD Bioscience,clone HIB19), CD161 (eBioscience, clone HP-3G10), CD127 (BioLegend, clone A019D5), and CD25 (Miltenyi, clone 4E3). Cells were sorted to exclude CD11c+ leukocytes and CD19+lymphocytes while including CD4+ T cells. CD25+, CD127- Tregs were sorted from the latter, and from the remaining CD4+ T cells, CD45RA- cells were sorted into CD161+ and CD161- fractions. 100,000 cells from each population were sorted directly into Trizol and frozen for RNA purification. RNA was later purified from these and sequenced by RNAseq on an Illumina HiSeq. Data were analyzed using GraphPad Prism software, version 8.4.2.

**CTLL2 overexpression of HPSE.** The murine HPSE transgene was co-expressed with an EGFP-Zeocin marker from a polycistronic message driven by the minimal EF1a promoter utilizing a self-cleaving P2A peptide. Lentiviral vectors were packaged in HEK 293T cells as previously described. Briefly, 10 µg of transfer plasmid was cotransfected with helper plasmids (10 µg of pCMV-Pol/Gag, 4 µg of pCMV-Rev, and 4 µg of pCMV-VSVG) into HEK 293T cells at 80 % confluence using Fugene 6 (Promega). Viral supernatant was harvested 48 hours post transfection, concentrated by ultracentrifugation, and stored at -80°C until use. Viral titers were determined by transduction of HT1080 cells and analyzed for EGFP expression by FACS. CTLL2 cells were transduced with viral supernatants at an MOI of 10. Protransduzin-A (Immundiagnostik AG) was added according to the manufacturer's instructions and transduction reactions were spinoculated at 1000 g for 90 minutes at 32°C in sealed tubes. An equal amount of fresh medium was added and cells were incubated in the presence of transduction complex for an additional 4 h at 37°C. The transduction medium was replaced with fresh medium and cells were cultured for 48 h before starting selection with 200 µg/ml Zeocin. After selection for 7 days > 99 % of cells were GFP positive by flow cytometry. Full vector sequences are available upon request.

**In vitro induction of HS-bound IL-2 signaling.** 96 well flat bottom tissue culture plates were pre-coated with subsequently anti-HS antibody (25 µg/ml) in PBS, 2.5 ug/ml HS in water, 10% BSA in PBS and 1000 IU/ml human recombinant IL2. Wells were washed 3 times with PBS in between each incubation step. CD4+ sorted cells were rested on ice in T cell media for 30 minutes before starting incubation by plating them on the pre-coated plates and gently spinning down for 30 seconds. After stimulation, cells were processed for phosphoflow staining as described above.

**Bone marrow chimeras.** Transfer of bone marrow cells from WT and $HPSE^{-/-}$ donor mice was performed as described previously[6]. In brief, recipient 6-8-week-old male CD45.1/CD45.2 F1 heterozygote recipient mice were lethally irradiated with 9Gy, rested for 24h and intravenously injected with 5 $x10^6$ bone marrow cells from WT and $HPSE^{-/-}$ mice at a 1:1 ratio. Chimerism was assessed 2 months after bone marrow transfer.

**Expansion and adoptive transfer of Treg.** Expansion of Treg was induced in WT and $HPSE^{-/-}$ -FOXP3.GFP mice by treatment with IL-2/anti-IL-2 antibody complex treatment as published before (Boyman et al., 2006). In short, complexes were prepared by incubating recombinant mouse IL-2 (eBioscience, 14-8021) and anti-IL-2 antibody (clone JES6-1, eBioscience, 16-7022) at a 1:5 ratio (w/w) for 30 min at 37 ˚C. After co-incubation, complexes were diluted with PBS and mice were administered 1 µg IL-2/5 µg anti-IL2 i.p. on 3 consecutive days. On day 5 after the first injection, spleens and lymph nodes were harvested and $CD4^+$/FOXP3.$GFP^+$ Treg were FACS sorted. For competitive adoptive transfer experiments, cells were washed, resuspended in PBS and mixed at a 1:1 ratio. A total of $2x10^6$ $CD25^+$cells were injected i.v. into 8-10-week-old male CD90.1 C57BL/6J mice. Distribution of donor Treg cells was analyzed 7 days after transfer. For treatment of EAE, IL-2/anti-IL-2 antibody complex expanded Treg were isolated using an EasySep™ mouse CD25 regulatory T cell positive selection kit (StemCell Technologies) according to manufacturer's directions. Cells were washed and resuspended in PBS and $1x10^6$ WT or $HPSE^{-/-}$ $CD25^+$ cells were transferred i.v. in 100 ul to 8-10-week-old female or male C57BL/6J mice one day before inducing EAE in the recipient mice as described above. Mice were monitored daily for clinical symptoms as follows: 0, no clinical disease; 1, tail weakness; 2, hindlimb weakness; 3, complete hindlimb paralysis; 4, hindlimb paralysis and some forelimb weakness; 5, moribund or dead.

**In vitro Treg suppression assay**. FACS sorted $CD4^+$/FOXP3.$GFP^+$ Treg were titrated into $1x10^5$ carboxyfluorescein succinimidyl ester (CFSE) labeled $CD4^+$/FOXP3.$GFP^-$ Tcon and co-cultured with $1x10^5$ irradiated $CD4^+$ T cell depleted splenocytes, together with 1ug/ml of soluble anti-CD3 (145-2C11, Biolegend). Proliferation of responder Tcon was assessed by flow cytometry after 72 hours of culture.

**Isolation of spinal cord infiltrating cells and flow cytometry.** For analysis of cell infiltrated into the central nervous system, pooled spinal cord tissue from 4-5 mice per experimental group, was homogenized using a dounce homogenizer. Cells were then isolated by density centrifugation using a 28% Percoll solution, layered with PBS. Cells were stained according to standard protocols with the following antibodies: CD3 (17A2), CD4 (GK1.5), CD25 (PC61), FOXP3 (FKJ.16a), Tbet (4B10), RORgT (B2D), GATA3 (TWAJ; all Biolegend) using staining reagents and protocols as per the manufacturer's instructions. Flow cytometry was performed on an LSRII (Becton Dickinson) in the Stanford Shared FACS Facility and data analysis was done using FlowJo (Treestar).

**In vitro CAR mRNA production, T cell transfection and transfer.** CAR HPSE and ix9Q mRNA were produced using T7 mScript™ Standard mRNA Production System from CellScript, following the manufacturer's protocols. T cells were transfected by electroporation using a Lonza nucleofector kit as per the manufacturer's protocol (Lonza). Briefly, $10x10^6$ cells were washed in PBS and mixed with CAR HPSE or ix9Q mRNA, as well as the Lonza reagent, in 100 ul and electroporated using an Amaxa system. Cells were cultured overnight to allow the chimeric receptor expression. Electroporated cells were then washed once more and injected into mice intravenously ($1x10^6$ to $1.5x10^6$ cells/mouse). For MOG-specific CAR Treg transfer, cells were incubated with a FITC-conjugated anti-MOG antibody in PBS for 30 minutes on ice and washed once before transfer.

**Statistical Analysis:** Statistical analysis was performed using GraphPad Prism software, version 8.4.2. A Kolmogorov-Smirnov test was used to verify normality of samples. In samples with Gaussian distribution, an unpaired t-test or 1-way ANOVA was used to determine significant differences between 2 or more groups, respectively, or a 2-way ANOVA was used to identify effects of multiple parameters. In samples without Gaussian distribution, a nonparametric Mann-Whitney test, or Friedman test for scoring over time, was used to determine significant differences. A p-value less than 0.05 was considered statistically significant.

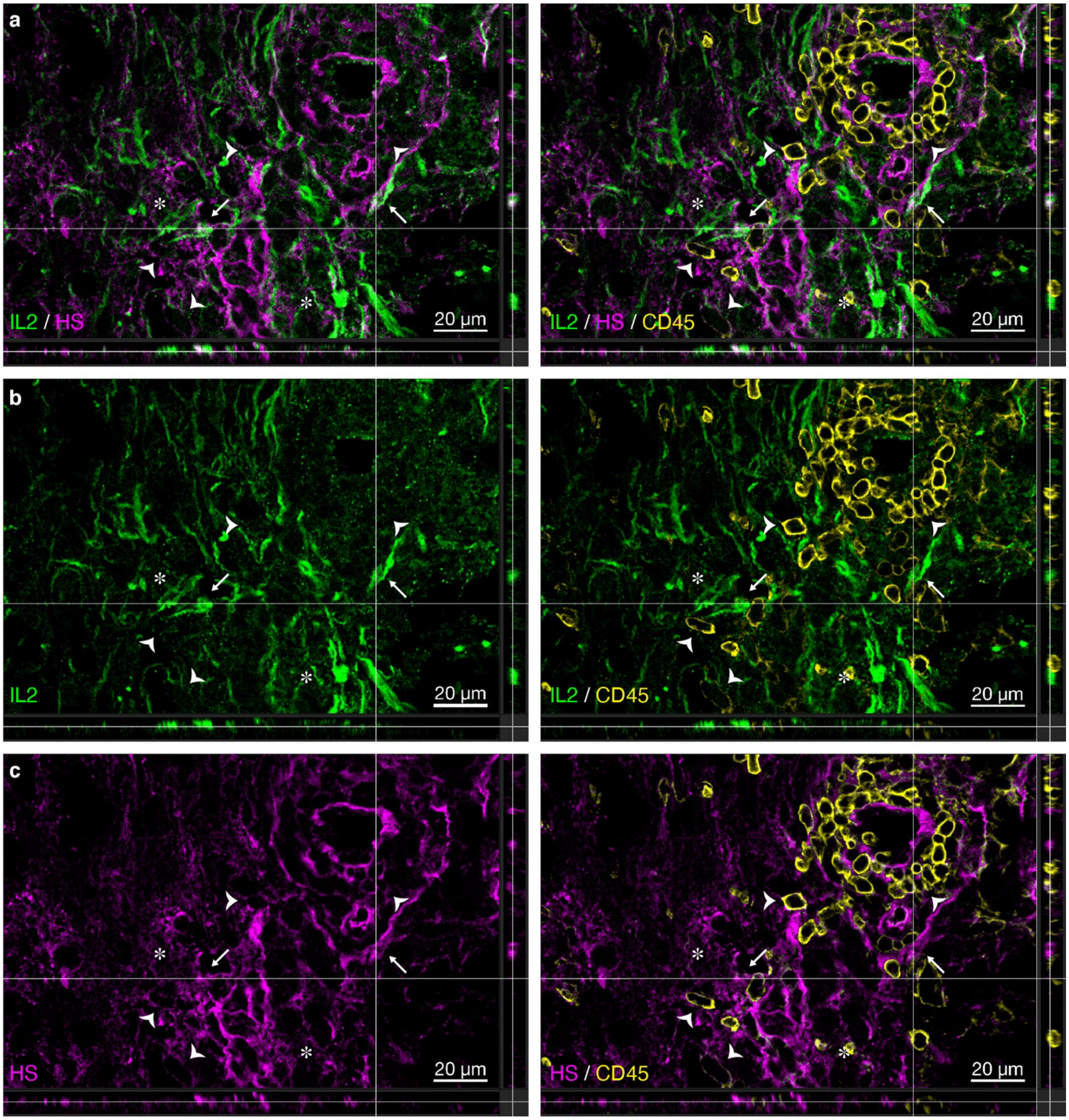


**Extended Data Figure 1. HS and IL2 colocalize in inflamed CNS tissue.** Orthogonal views of z-stacks of EAE spinal cord tissue (29 dpi) stained for IL-2 (green), HS (magenta) and CD45 (yellow). Shown are the overlay of IL-2 and HS staining (**a**, overlay with appear white/grey), as well as individual IL-2 (**b**) and HS (**c**) channels. Arrows indicate overlap of IL-2 and HS staining on cellular structures, arrowheads indicate HS staining associated with CD45 cells and asterisks indicate diffuse IL-2 and HS staining.

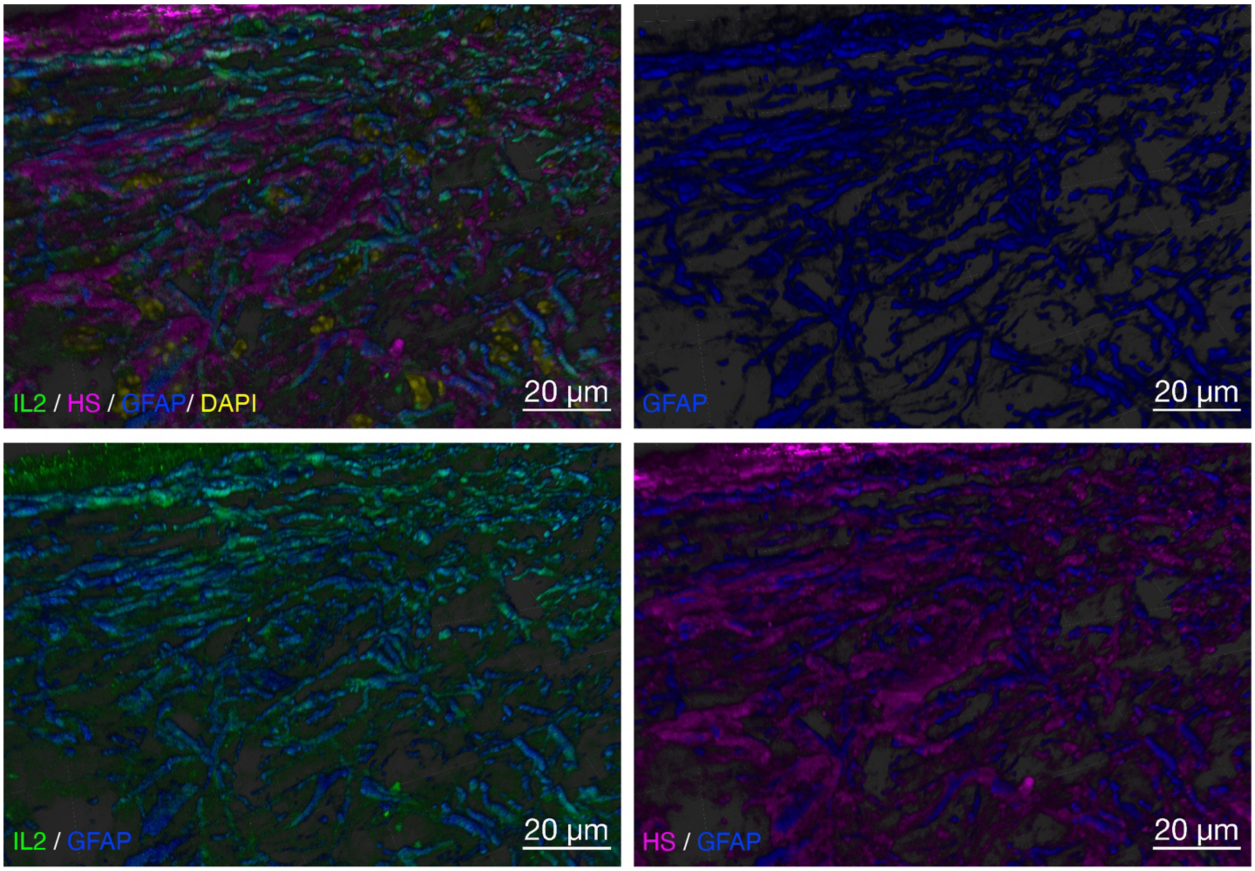


**Extended Data Figure 2. Immunohistochemical analysis of GFAP, HS and IL2 in EAE CNS tissue.** Imaris 3D surface rendering of EAE spinal cord tissue stained for GFAP (blue), IL-2 (green) and HS (magenta), showing association of both IL-2 and HS staining with GFAP positive structures. DAPI counterstain for nuclei is shown in yellow.

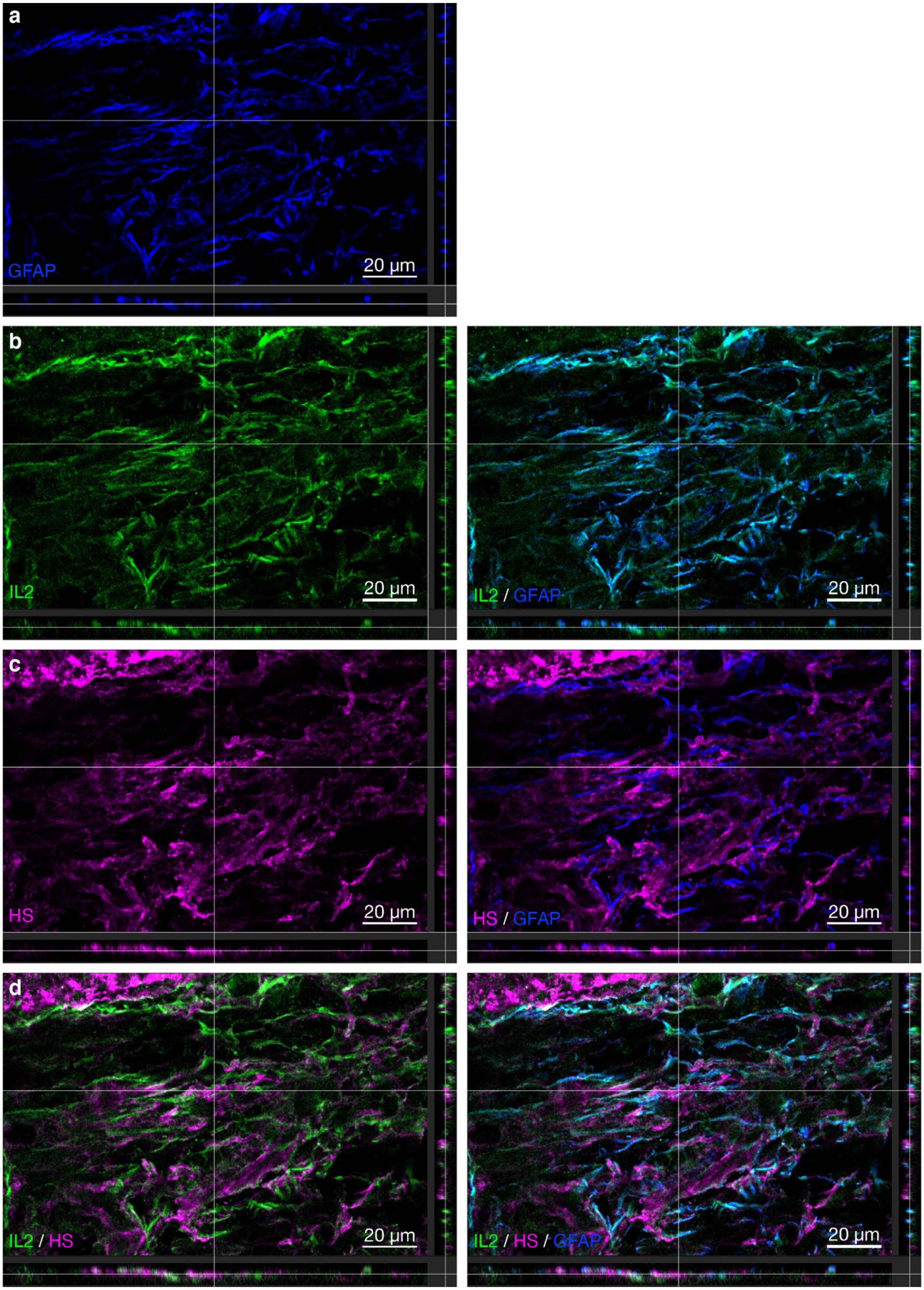
a
GFAP
20 µm
b
IL2
20 µm
IL2 / GFAP
20 µm
c
HS
20 µm
HS / GFAP
20 µm
d
IL2 / HS
20 µm
IL2 / HS / GFAP
20 µm

1

**Figure S3. IL-2 and HS are associated with reactive astrocytes.** Orthogonal views of z-stacks of EAE spinal cord tissue stained for GFAP (blue), IL-2 (green) and HS (magenta). Shown are individual GFAP (A), IL-2 (B) and HS (C) channels, as well as the overlay of IL-2 and HS staining (D, overlay with appear white/grey), alone and in conjunction with GFAP staining.

**a**

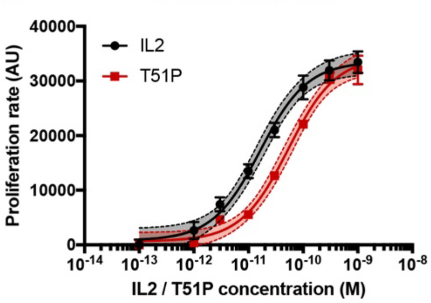


| | EC50 (M) | 95% CI (M) | $R^2$ (non-linear regression) |
|---|---|---|---|
| **IL2** | 1.622e-011 | 1.077e-011 to 2.443e-011 | 0.9803 |
| **T51P** | 5.353e-011 | 3.775e-011 to 7.589e-011 | 0.9846 |

**b**

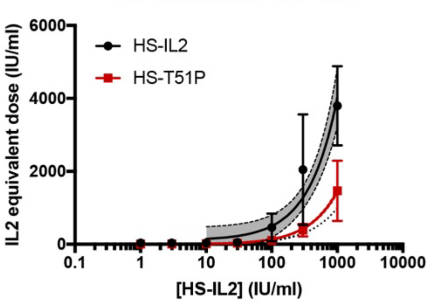


| HS-bound concentration (IU/ml) | Equivalent soluble concentration (IU/ml) [fold increase] | |
|---|---|---|
| | **IL2** | **T51P** |
| 1 | 113 [113 x] | - * [-] |
| 3 | 120 [40 x] | - * [-] |
| 10 | 148 [14.8 x] | 6 [0.6 x] |
| 30 | 226 [7.53 x] | 35 [1.1 x] |
| 100 | 500 [5 x] | 138 [1.38x] |
| 300 | 1281 [4.27 x] | 430 [1.43 x] |
| 1000 | 4018 [4.018 x] | 1452 [1.452 x] |
| $R^2$ (linear regression) | 0.9486 | 0.9979 |

**c**

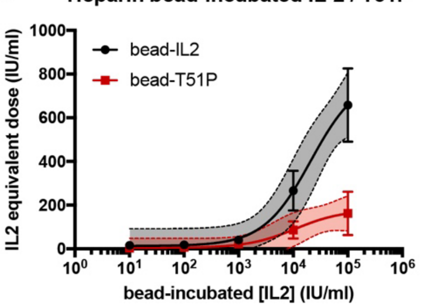


| Bead-incubated concentration (IU/ml) | Equivalent soluble concentration (IU/ml) | |
|---|---|---|
| | **IL2** | **T51P** |
| $1*10^1$ | 16 | 3 |
| $1*10^2$ | 17 | 6 |
| $1*10^3$ | 41 | 23 |
| $1*10^4$ | 266 | 86 |
| $1*10^5$ | 658 | 162 |
| $R^2$ (non-linear regression) | 0.821 | 0.6482 |

**Extended Data Figure 4. Quantification of effective doses of HS-bound and heparin-coated bead-bound IL-2 and T51P-IL-2.** Shown are mean +/- SEM for experimental values, best-fit curves for each analysis and 95% confidence areas (shaded areas). **a,** Non-linear regression analysis (left; variable slope, 3 parameters) and calculated EC50 values (right) of proliferation of CTLL2 cells induced by soluble IL-2 and T51P-IL-2. **b,** Linear regression analysis (left) and extrapolated effective doses of HS-bound IL-2 and T51P-IL-2 assessed by induction of proliferation of CTLL2 cells (n=2-4 experiments). The dotted line depicts a theoretical curve of equal effectiveness (comparing bound IL-2 to soluble IL-2). Table (right) shows extrapolated equivalent doses delivered by HS-bound IL-2 concentrations, depicting which dose of soluble IL-2, or T51P-IL-2, elicits a similar proliferative response as the dose of HS-bound IL-2, or T51P-IL-2, shown in the left column. Fold increase of proliferation rate over the rate induced by the same unbound cytokine is indicated as well. * below detection. **c,** Non-linear regression analysis (left; variable slope, 4 parameters) and extrapolated effective doses of heparin-coated bead-bound IL-2 and T51P-IL-2, assessed by induction of proliferation of CTLL2 cells (n=3 experiments). Table (right) shows extrapolated equivalent doses IL-2 delivered by heparin coated beads pre-incubated with various concentrations of IL-2, depicting which concentration of soluble IL-2, or T51-P, elicits a similar proliferative response as heparin beads pre-incubated with the concentration of IL-2, or T51P-IL-2, shown in the left column.

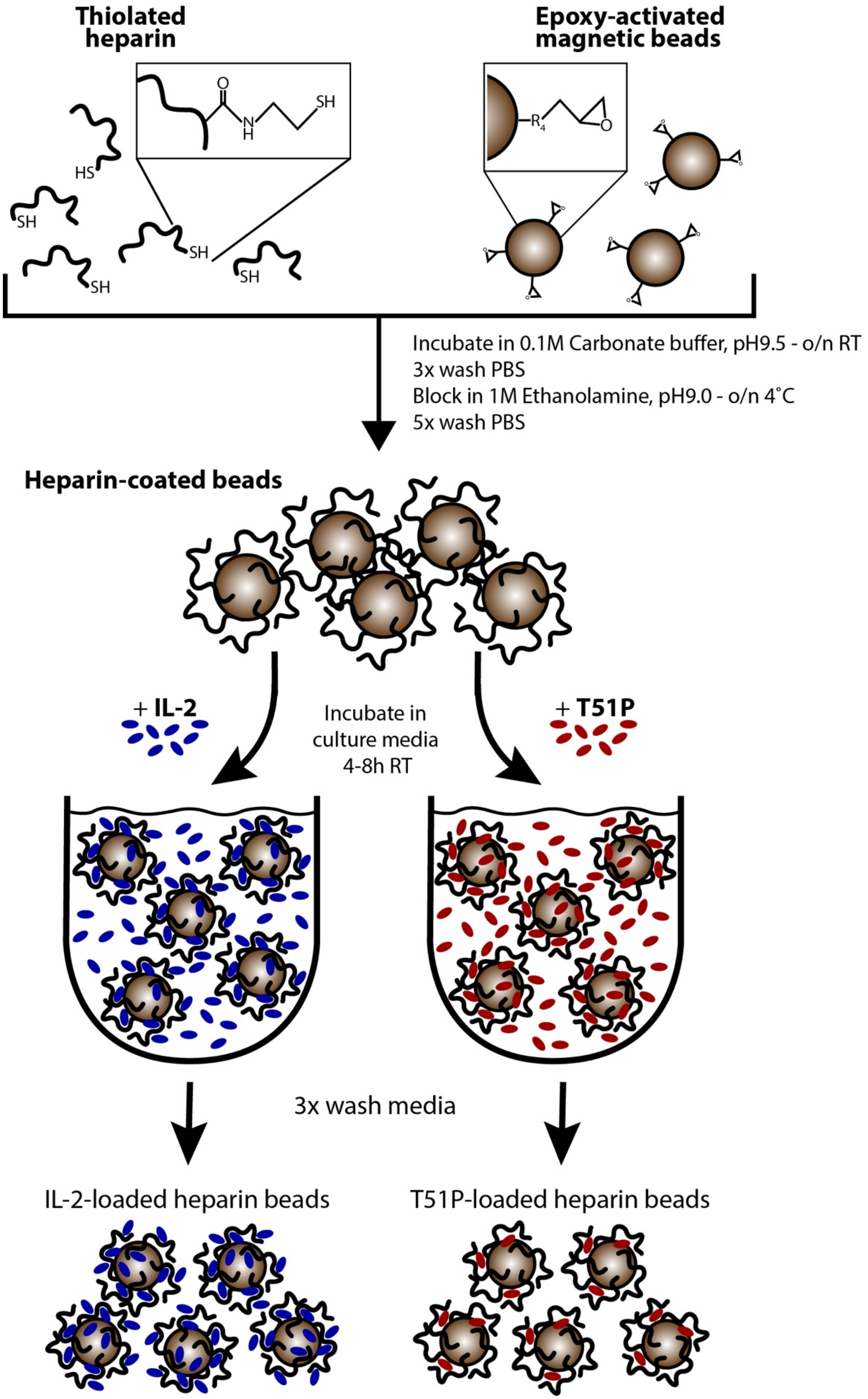
Thiolated heparin
O
N
H
SH
HS
SH
SH
SH
SH
Epoxy-activated magnetic beads
R4
O
Incubate in 0.1M Carbonate buffer, pH9.5 - o/n RT
3x wash PBS
Block in 1M Ethanolamine, pH9.0 - o/n 4˚C
5x wash PBS
Heparin-coated beads
+ IL-2
Incubate in culture media 4-8h RT
+ T51P
3x wash media
IL-2-loaded heparin beads
T51P-loaded heparin beads

1

**Extended Data Figure 5. Generation of IL-2-loaded heparin-coated beads.** Schematic overview of generation of IL-2-loaded, heparin-coated magnetic beads. Thiolated heparin is coupled to 1 μm epoxy-activated magnetic beads to yield heparin-coated beads. These beads are then incubated with IL-2 or T51P-IL-2 in cell culture media (RPMI containing 10% FCS and penicillin/streptomycin). After washing, these IL-2 or T51P-loaded beads are used to deliver IL-2 to cells in culture.

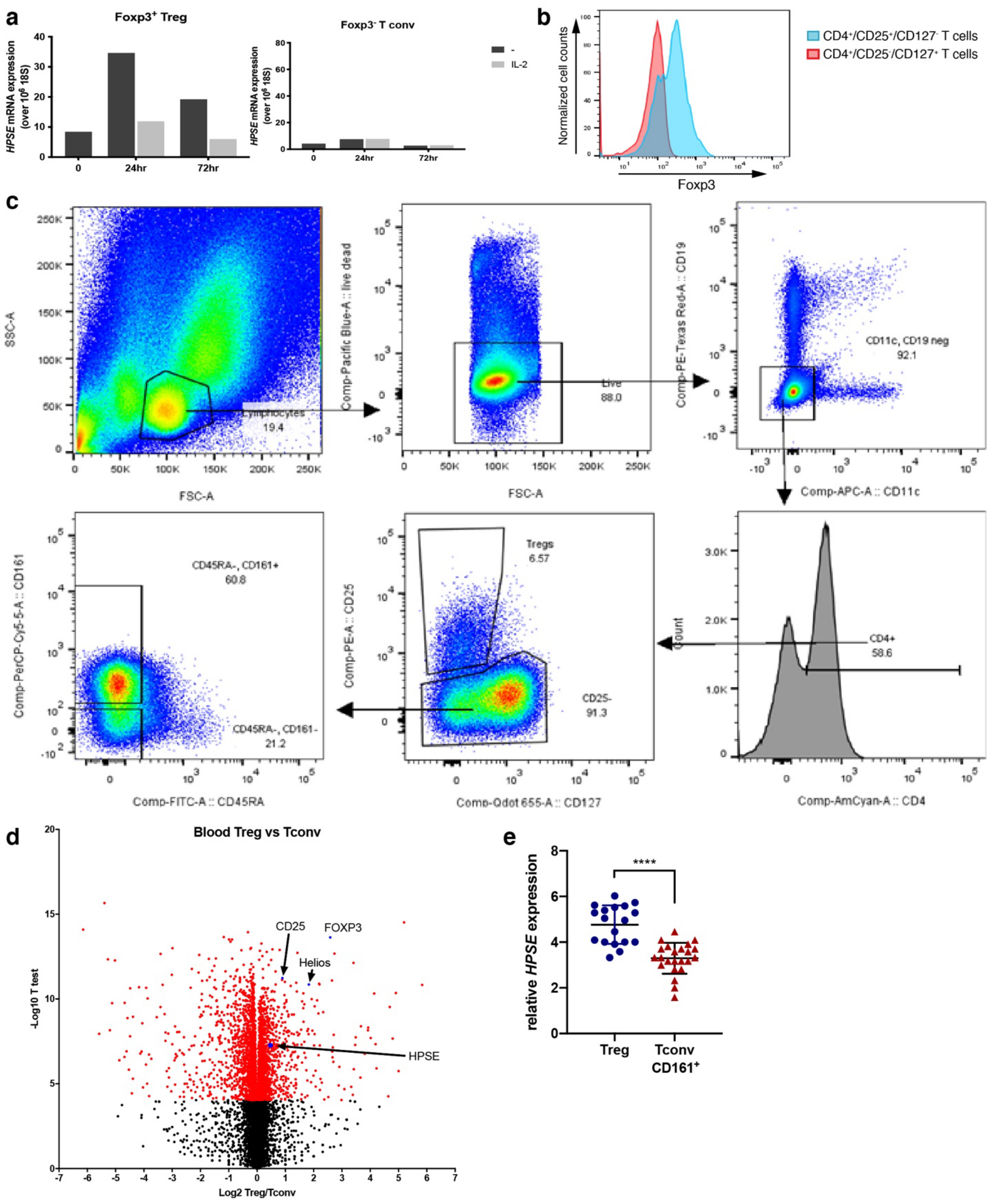


**Extended Data Figure 6. Treg express higher levels of HPSE than Tconv. a,** Upregulation of HPSE in Treg is regulated by IL-2: qPCR quantitation of *Hpse* mRNA expression in FACS sorted murine Foxp3+ Treg and Foxp3- Tconv after *in vitro* activation with aCD3/aCD28, either alone, or in the presence of IL-2 (200 IU/ml). mRNA expression was normalized by *18S* mRNA expression. **b,** Confirmation of Foxp3 expression in FACS sorted human Treg**.** Foxp3 protein expression in FACS sorted and *in vitro* activated human CD4+/CD25+/CD127- Treg (blue histogram) and CD4+/CD25-/CD127+ Tconv (red histogram). **c,** Human Treg express higher levels of HPSE than Tconv: Gating scheme showing the Treg and Teff populations that were isolated from human blood and colon samples and analyzed for gene expression. **d,** Volcano plots depicting relative expression of genes in Treg over Teff in human blood. **e,** Relative *HPSE* mRNA expression in FACS sorted human Treg and $CD161^+$ Tconv isolated from surgically resected colons. Shown are mean relative HPSE expression +/- SD **** $p < 0.0001$, one-way ANOVA.

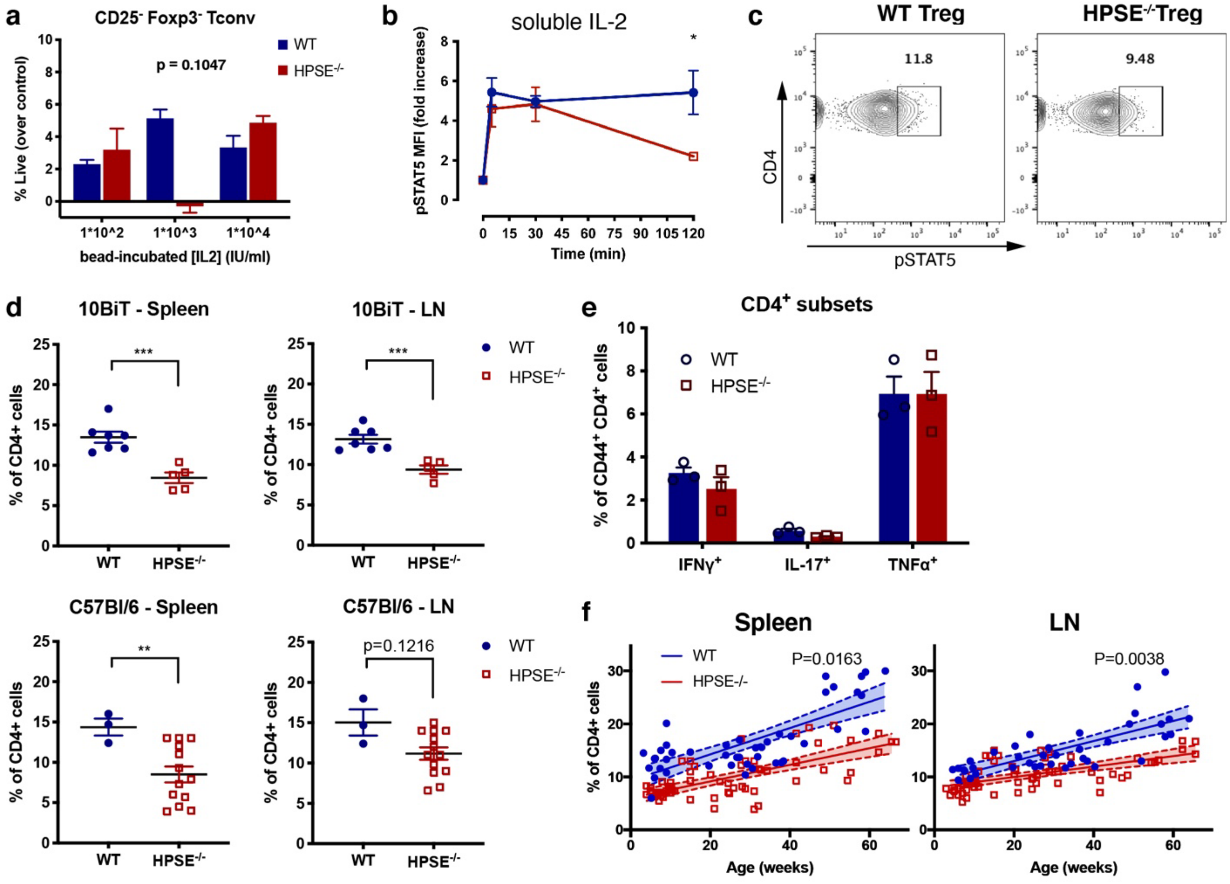


**Extended Data Figure 7. HPSE expression supports Foxp3+ Treg access to IL-2 and their homeostasis *in vitro* and *in vivo*. a,** Viability of WT and HPSE$^{-/-}$ CD25$^{-}$ Foxp3$^{-}$ Tconv among CD4$^{+}$ T cells cultured in the presence of heparin-coated beads pre-incubated with increasing concentrations of IL-2. Viability was measured by flow cytometry 24 hr after start of culture and is shown as percent viable cells corrected for baseline viability of cell cultured in media alone. Shown is a representative of 4 independent experiments (mean + SEM of triplicate samples); p-value depicts variation due to the genotype (WT vs. HPSE$^{-/-}$), determined by two-way ANOVA. **b,** Stat5 phosphorylation in WT and HPSE$^{-/-}$ Foxp3$^{+}$ Treg after stimulation with soluble IL-2. * $p < 0.05$, two-way ANOVA with Sidak's multiple comparison correction. **c,** Representative plots of the percentage of pStat5+ cells among CD4$^{+}$ Foxp3$^{+}$ Treg isolated from naïve WT and HPSE$^{-/-}$ spleen tissue. **d,** Percentage of Foxp3+ Treg among CD4+ T cells in the spleens (lefts panel) and inguinal lymph nodes (right panel) of adult (3 to 6-month-old) WT and HPSE-/- mice on the 10BiT background (top panels) C57Bl/6 background (lower panels). ** $p < 0.01$, *** $p<0.001$, two-tailed t-test. **e,** IFN-γ, IL-17 and TNF-α-producing CD4+ T cell subsets in mesenteric lymph nodes of WT and HPSE$^{-/-}$ mice; $n = 3$ mice. **f,** Linear regression analysis of Foxp3+ Treg frequencies among CD4+ T cells in the spleens (left panel) and inguinal lymph nodes (right panel) of WT and HPSE$^{-/-}$ mice during aging. p-value shown comparing slopes.

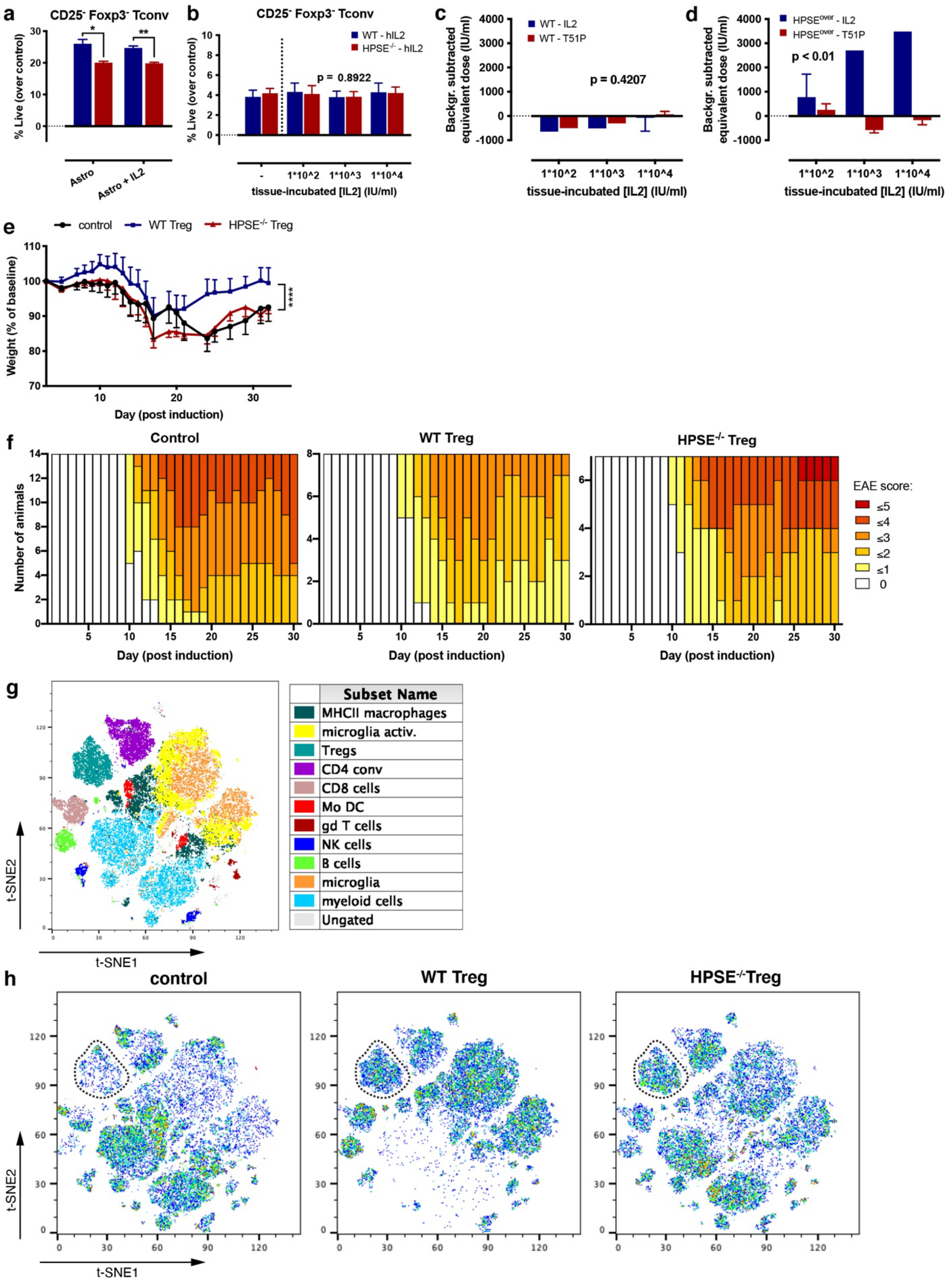

a
CD25- Foxp3- Tconv
% Live (over control)
Astro
Astro + IL2
b
CD25- Foxp3- Tconv
WT - hIL2
HPSE-/- - hIL2
p = 0.8922
% Live (over control)
tissue-incubated [IL2] (IU/ml)
c
WT - IL2
WT - T51P
p = 0.4207
Backgr. subtracted equivalent dose (IU/ml)
tissue-incubated [IL2] (IU/ml)
d
HPSEover - IL2
HPSEover - T51P
p < 0.01
Backgr. subtracted equivalent dose (IU/ml)
tissue-incubated [IL2] (IU/ml)
e
control
WT Treg
HPSE-/- Treg
Weight (% of baseline)
Day (post induction)
f
Control
WT Treg
HPSE-/- Treg
Number of animals
Day (post induction)
EAE score:
≤5
≤4
≤3
≤2
≤1
0
g
Subset Name
MHCII macrophages
microglia activ.
Tregs
CD4 conv
CD8 cells
Mo DC
gd T cells
NK cells
B cells
microglia
myeloid cells
Ungated
t-SNE1
t-SNE2
h
control
WT Treg
HPSE-/-Treg
t-SNE1
t-SNE2


1

**Extended Data Figure 8. HPSE expression supports Foxp3$^+$ Treg access to tissue-bound IL-2 and their function *in vivo.*** **a** and **b,** Viability of WT and HPSE$^{-/-}$ CD25$^-$ Foxp3$^-$ Tconv among CD4$^+$ T cells cultured in the presence of U87-MG cells (**a**), or mouse spinal cord tissue (**b**) pre-incubated with IL-2. Viability was measured by flow cytometry 24 hr after start of culture and is shown as percent viable cells corrected for baseline viability of cell cultured in media alone. Shown are representatives of 4 (**a**) or 3 (**b**) independent experiments (mean + SEM of triplicate samples); (**a,** * p < 0.05, ** p<0.01, two-tailed t-test with Holm-Sidak multiple comparison correction; **b,** p-values depict variation due to the genotype (WT vs. HPSE$^{-/-}$), determined by two-way ANOVA. **c** and **d,** Proliferation of (**c**) WT and (**d**) HPSE overexpressing CTLL2 cells in response to freshly isolated and irradiated mouse spinal cord tissue that was pre-incubated with IL-2 or T51P-IL-2. Proliferation rate is depicted as equivalent dose of soluble IL2 or T51P-IL-2 at which a similar proliferative response is elicited, corrected for background proliferation induced by spinal cord tissue alone. Shown are mean + SEM of 1-3 wells per sample, p-value depict variation due to the cytokine that the beads were incubated with (IL-2 vs. T51P-IL-2), determined by two-way ANOVA. **e,** Weights of EAE -afflicted animals that received WT or HPSE$^{-/-}$ Foxp3$^+$ Treg 1 day prior to induction of EAE. Average weight +/- SEM are shown as percentage of original weight at d0. **** p<0.0001, two-way ANOVA with Tukey's multiple comparison correction between WT Treg and control, and WT Treg en HPSE$^{-/-}$ Treg. Shown is a representative of 2 independent experiments. **f,** Distribution of disease scores among the treatment groups. The number of animals with indicated color-coded severity scores are shown. **g,** Unsupervised clustering t-distributed stochastic neighbor embedding (tSNE) plot of flow cytometry (protein expression) analysis of cells in spinal cord tissue at peak of disease (d21-25), identifying immune cell subsets. **h,** tSNE plots for animals that did not receive Treg (control), and animals that received WT and HPSE-/- Foxp3+ Treg 1 day prior to induction of EAE. The cell cluster identified as Treg is indicated. tSNE analysis was performed in FLowJo.

**Extended Data Table 1.**

**EAE statistics.** Shown is a representative of 2 independent experiments.

| Experimental group | Incidence (%) | Disease onset† (day) | Peak severity† (score) | Average severity† d12-30 (score) |
|---|---|---|---|---|
| **Control** | 100% (14/14) | 10.8 ± 0.4 | 3.5 ± 0.2 | 2.5 ± 0.2 |
| **WT Treg** | 100% (8/8) | 11.0 ± 0.3 | 2.6 ± 0.2* | 1.7 ± 0.2* |
| **HPSE$^{-/-}$ Treg** | 100% (7/7) | 11.0 ± 0.3 | 3.5 ± 0.3**** | 2.6 ± 0.4 |

† Listed are mean values +/- SEM per group

* p<0.05, **** p<0.0001, Kruskal-Wallis test comparing WT Treg vs. control or HPSE$^{-/-}$ Treg vs. WT Treg.

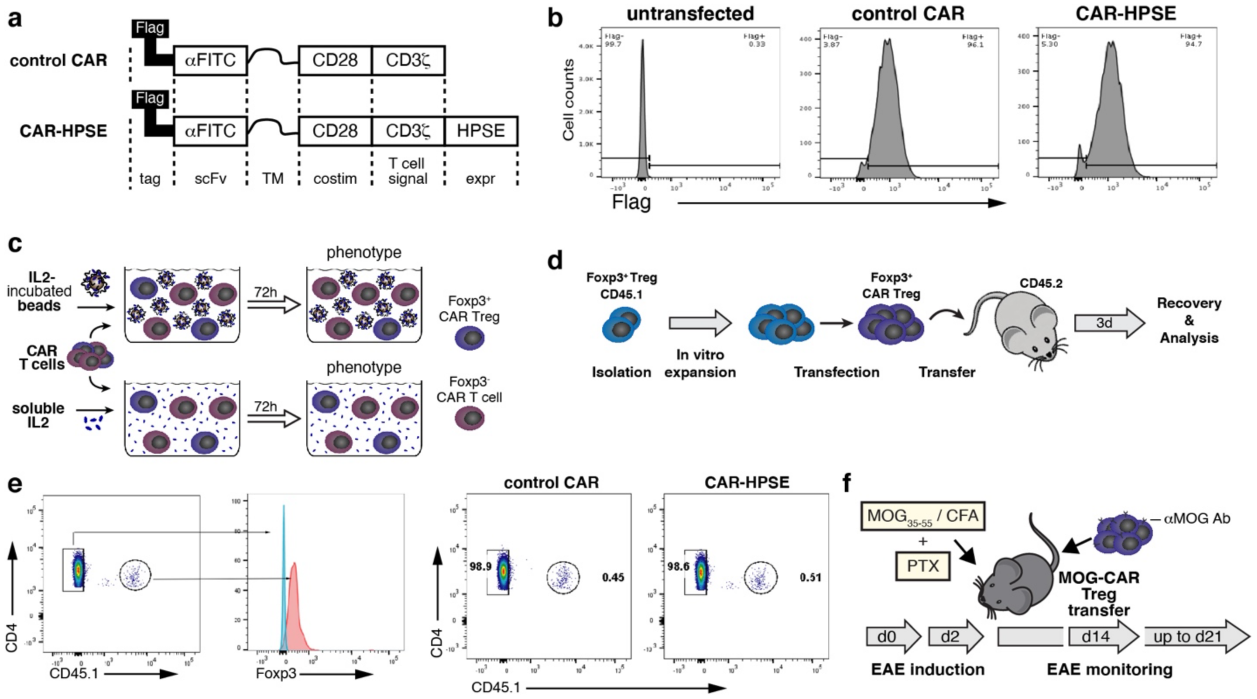


**Extended Data Figure 9. HPSE overexpression enhances phenotypic stability of CAR-Treg.** **a,** Schematic overview of the chimeric antigen receptor (CAR) constructs used to transfect into Treg. Abbreviations: scFv - single-chain variable region; TM - transmembrane domain; costim - costimulatory signaling domain; T cell signal - T cell receptor signaling domain; expr - expression module. **b,** Flow cytometry histograms depicting the transfection efficiency of the CAR construct used. Efficiency was determined by staining for the FLAG tag present in the constructs. Shown are representative plots of 3 independent experiments. **c,** Schematic overview of the assay used to assess in vitro stability of CAR-Treg (either control, or overexpressing HPSE) induced by heparin-coated beads pre-incubated with IL-2, or soluble IL-2. **d,** Schematic overview of the experiment designed to assess the *in vivo* stability of transferred control and HPSE CAR Treg (Foxp3+). **e,** Identification strategy (left panels) and frequency (right panels) of control and HPSE overexpressing CAR-Tregs (CD45.1+) among CD4+ T cells in lymphoid tissue, 3 days after adoptive transfer of these cells into C57Bl/6 recipients (CD45.2+). **f,** Schematic overview of the experiment designed to assess the capacity of transferred control and HPSE-overexpressing MOG-specific CAR Treg (Foxp3+) to suppress EAE.

### Extended Data References
